# Generation and long-term expansion of human pancreatic islet organoids *in vitro*

**DOI:** 10.64898/2026.04.15.718643

**Authors:** Wenqian Song, Chunye Liu, Shusen Wang, Daisong Wang, Yishu Xu, Shubo Yuan, Jiali Chang, Boya Zhang, Xu Han, Hongxing Fu, Haili Bao, Aijing Shan, Di Zheng, Wenquan Wang, Yanan Cao, Weiqiong Gu, Jiqiu Wang, Liang Liu, Shaohua Song, Qing Cissy Yu, Yi Arial Zeng

## Abstract

The scarcity of expandable, functional human islet cells remains a major barrier to diabetes therapy. Here, we identify PROCR+ cells within adult human islets and establish a defined culture system to generate pancreatic islet organoids. These organoids self-organize into α-, β-, δ-, and PP cells at near-native ratios, exhibit regulated insulin and glucagon secretion, and support exponential in vitro expansion. Single-cell transcriptomics reveals a unique progenitor-like cell population that is transcriptionally primed for endocrine differentiation but shares molecular features with fetal trunk cells and endocrine progenitors. When transplanted, the organoids rapidly ameliorate hyperglycemia in diabetic mice. Importantly, in a non-human primate model, intraportal transplantation of these organoids reduced exogenous insulin requirements, restored glucose-stimulated C-peptide secretion, and achieved sustained glycemic control-representing a critical step toward clinical translation. This study provides a strategy for expanding human islet organoids, offering a scalable platform for diabetes treatment, disease modeling, and regenerative medicine.

## Introduction

The islets of *Langerhans*, the endocrine compartment of the pancreas, regulate whole-body glucose homeostasis through the secretion of hormones like insulin (β cells), glucagon (α cells), somatostatin (δ cells), and pancreatic polypeptide (PP/γ cells) in response to nutrient levels^1–5^. Insulin and glucagon are the primary outputs, with insulin secretion stimulated post-feeding and glucagon released during fasting to prevent hypoglycemia. This balance is maintained with remarkable precision in healthy islets^6–8^.

Pancreatic islets are not mere aggregates of endocrine cells but rather highly organized mini-organs^6,9,10^. Unlike murine islets, human islets exhibit significant variability in cellular composition, with β cells comprising 30%-75%, α cells 10%-40%, and δ and PP cells each 1%-10% of the islet population^1,3,11–13^. Intra-islet interactions, particularly feedforward signaling between α and β cells, are essential for glucose-stimulated insulin secretion (GSIS), while feedback mechanisms involving δ cells prevent excessive insulin release, safeguarding against hypoglycemia^6,14–18^.

Diabetes mellitus arises when pancreatic β cells fail to secrete sufficient insulin to meet physiological demands, leading to hyperglycemia^19^. Transplantations of whole pancreas or isolated islets from cadaveric donors are feasible treatment for restoring glucose homeostasis by replenishing the functional β cell pool^20–22^. However, the scarcity of donor organs has spurred the search for strategies to generate β cells in vitro. Although *in vitro* expansion of islets represents an ideal therapeutic approach, the limited proliferative capacity of mature endocrine cells poses a significant obstacle^23,24^.

Pancreatic islet development and regeneration involve layers of regenerative mechanisms^25^, including β cell replication^26^and islet-resident progenitor cell differentiation^27,28^ during homeostasis. Upon injury, ductal cells reactivate differentiation programs^29–31^, and acinar cells transdifferentiate to generate new islet cells^32–34^. The key challenge is leveraging these mechanisms to generate new β cells, or ideally, regenerate entire islets in vitro.

Significant progress has been made in developing pancreatic organoids from both fetal and adult mouse^35–38^ and human^39–42^ pancreas. Notably, exocrine ductal cells have demonstrated the ability to form organoids that can be expanded and maintained in vitro for extended periods. These ductal organoids have the potential to differentiate into both exocrine and endocrine cell types, suggesting that there may exist a subpopulation of duct cells with inherent adult progenitor capacity or the ability to acquire cellular plasticity under specific conditions. However, they predominantly express high levels of ductal genes and show limited efficiency in differentiating into endocrine islet cells^38,43,44^. Thus, efficient generation of endocrine organoids from primary tissues remains a significant challenge.

Our previous study has identified a population of Protein C Receptor (Procr)-expressing resident progenitor cells within mouse pancreatic islets^28^. Procr is recognized as a marker of adult stem cells across various tissues^45–54^. Procr+ stem cells exhibit enhanced resilience to cytotoxic stress^55^, and the ability to endure and proliferate in vitro ^28,49,50,56,57^. In mouse islets, Procr+ progenitors constitute ∼1% of the islet cell population and lack expression of known islet hormones. Lineage tracing has shown their capacity to generate all four endocrine cell types in vivo, and they can be expanded long-term in vitro to form islet-like organoids containing all relevant endocrine cell types^28^.

Building on these findings, the current study investigates the presence of PROCR+ cells in the human pancreas and explores their potential for generating human pancreatic islet organoids in vitro.

## Results

### Visualization of PROCR+ cells in human Pancreas islets

To investigate the presence of PROCR+ cells within human islets, we performed immunostaining on human pancreas tissue sections. We identified a small population of PROCR+ cells within the islets. Notably, these cells were hormone-negative, showing no detectable staining of insulin (INS), glucagon (GCG), somatostatin (SST), and pancreatic polypeptide-secreting (PPY), similar to the Procr+ cells observed in mouse islets (Figure S1A).

We further analyzed freshly isolated islets using whole-mount staining. This approach confirmed that the PROCR+ cells located within the islets were hormone-negative (Figures 1A, 1B). In instances where isolated islets were contaminated with attached exocrine tissues, we observed that these exocrine tissues contained SOX9+ ductal cells. In contrast, within the islets, PROCR+ cells were SOX9- and positive for the pancreatic marker PDX1 (Figure 1C).

**Figure 1.**
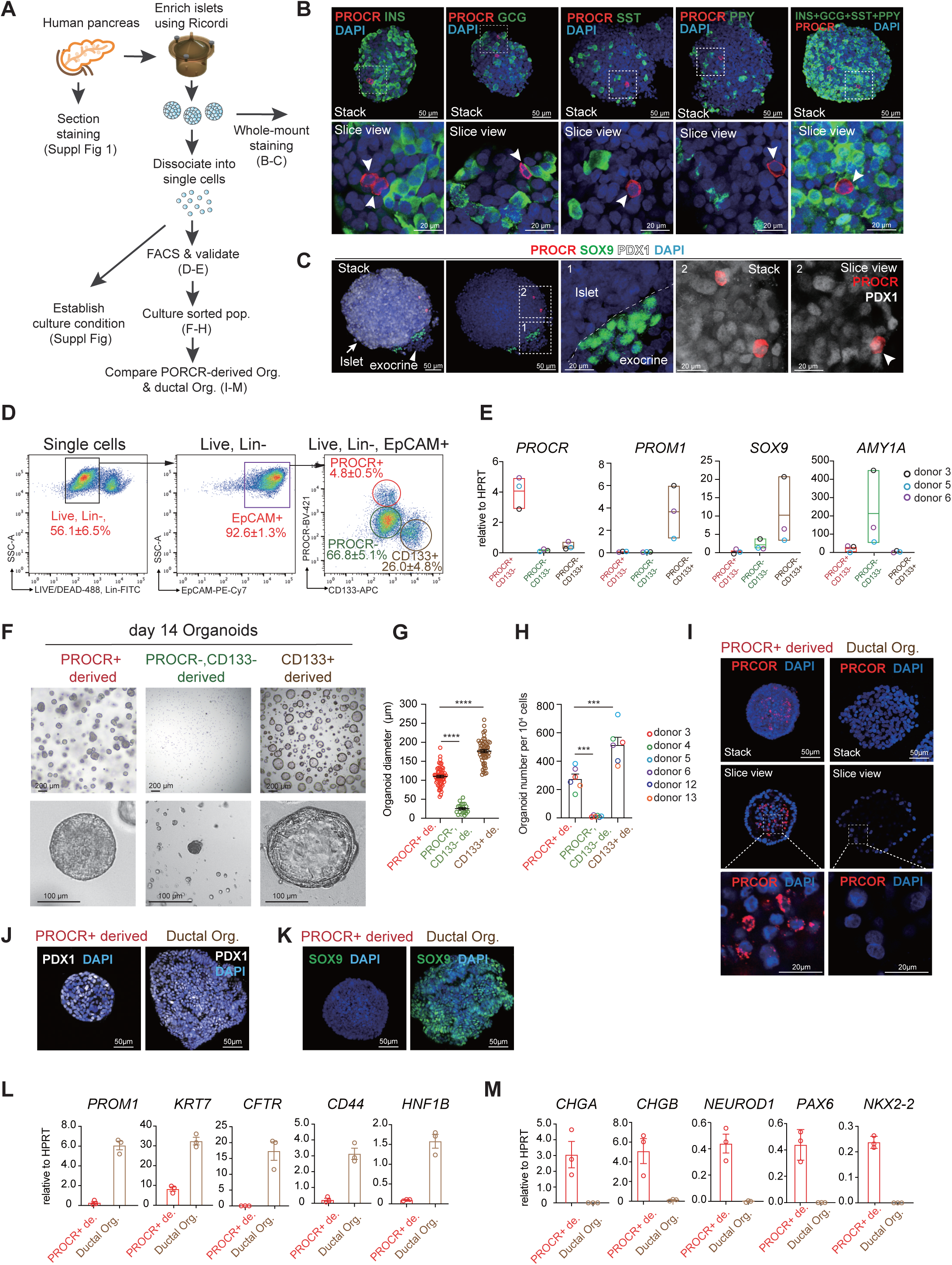
Human pancreatic PROCR+ cells generate unique organoids distinct from ductal organoids. (A) Experimental diagram. Pancreatic islets are isolated and enriched by perfusion, enzymatic digestion, and density gradient centrifugation as previously described. Freshly isolated islets are used for whole-mount immunofluorescence staining. The dissociated islet single cells are used for cell culture or cell sorting. (B) Representative confocal images of human pancreatic islets (Donor #13) immunostained for PROCR (red) and endocrine markers (green). Upper panels showing Z-stack views, and lower panels display magnified slice views (single-plane). Arrows denote PROCR+ cells lacking co-localization with β cells (INS+), α cells (GCG+), δ cells (SST+), or PP cells (PPY+). Scale bar, 50 µm (upper panels) and 20 µm (lower panels). Data representative of islets from n = 3 independent donors (see Table S1). (C) Representative confocal images of human pancreatic islets (Donor #12) immunostained for PROCR (red), ductal maker SOX9 (green), and pancreatic marker PDX1 (white). Left: Z-stack projection (scale bar, 50 µm). Middle: magnified regions 1 and 2 (scale bar, 20 µm). Right: Orthogonal slice view of region 2 (scale bar, 20 µm). Arrowheads denote PROCR+, PDX1+ cell within the islet. Data representative of islets from n = 3 independent donors (see Table S1). (D) Representative FACS analysis of PROCR+ cells in primary human pancreatic islet cells. The dissociated islet single cells were stained with Violet Live/Dead dye to exclude dead cells, selected for Lin- (CD45- and CD31-) to exclude blood and endothelial lineage, and selected for EpCAM+ to enrich epithelial cells. CD133 is used to exclude duct cells. The percentage of PROCR+ cells in live, Lin-, EpCAM+ cell compartment was analyzed. The FACS plot presents data from donor #3, while the quantified percentages reflect the averaged data (mean ± SEM) across 10 independent donors. Results for each individual donor are presented in Figure S2. (E) RT-qPCR analysis of different markers expression in FACS sorted three populations. Data were pooled from n=3 donors (see Table S1), presented as floating bars spanning minimal to maximum. Each dot represents an organoid line, with data generated from three technical replicates. (F-G) PROCR+ or PROCR-, CD133- or CD133+ cells were FACS sorted (from Live, Lin-, EpCAM+ compartment) and cultured. Representative bright-field images of colony formation at culture day 14 are shown (donor #13) (F). Quantification of organoid sizes for each cultured group are shown (G). Scale bar, 200 µm in upper panels and 100 µm in lower panels (F). Data were pooled from n=6 donors (see Table S1), presented as mean ± SEM. One-way ANOVA with Tukey’s test is used for the comparison of multiple groups. **** p<0.0001. (H) Organoid numbers derived from cultured PROCR+ or PROCR-, CD133- or CD133+ cells are shown. Data are from n=6 donors (represented by distinct circle colors) (see Table S1), presented as mean ± SEM. One-way ANOVA with Tukey’s test is used for the comparison of multiple groups. *** p<0.001. (I-K) Representative confocal images of PROCR+ cell derived and CD133+ cell derived organoids from Donor #12, immunostained for PROCR (I), PDX1 (J), and SOX9 (K). Scale bar, 50 µm. Scale bar, 20 µm (magnified regions in I). Experiment repeated with samples from n = 3 donors (see Table S1). (L-M) RT-qPCR analysis of duct maker (L) and endocrine marker (M) expression in PROCR+ cell derived and CD133+ cell derived organoids. Experiment repeated with samples from n=3 donors (see Table S1). Each symbol represents an organoid line, with data generated from three technical replicates.

### A distinct type of human pancreatic organoids from PROCR+ cells

Next, we sought to isolate live PROCR+ islet cells. Human pancreas tissue samples were obtained from deceased organ donors and subjected to enzymatic digestion followed by density gradient centrifugation to purify islets, adapting a method originally described by Ricordi et al.^58,59^. This islet-enriched fraction underwent single-cell dissociation and Fluorescence-activated cell sorting (FACS) analysis (Figures 1A, 1D). Live cells were initially selected, and blood and endothelial lineage cells were depleted (Live, Lin-). The epithelial compartment (EpCAM+) was then gated, and PROCR and CD133 expression were analyzed. We found that PROCR+ cells constituted approximately 4.8 ±0.5% of total EpCAM+ islet cells (n=10 donors) (Figure 1D, Figure S2A-B, Table S1).

The percentage of PROCR+ cells identified by FACS analysis appeared higher than that observed through immunofluorescence staining of tissue sections. We noted a significant proportion of dying cells (∼ 43.9%), resulting from the islet digestion and enrichment process (Figure 1D, Figure S2). Given the known resilience of PROCR+ cells to cytotoxic stress^55^, we hypothesized that they may better endure the isolation procedures than mature islet cells, potentially leading to an overrepresentation of PROCR+ cells in the FACS analysis. Due to the propensity of pancreatic duct cells to proliferate in culture^38,60–63^, we also analyzed CD133+ duct cells (Figure 1D, Figure S2) to exclude potential ductal contamination from the islet cell-enriched population. RT-qPCR analysis of *PROCR* and markers of pancreatic duct cells, *SOX9* and *PROM1*, confirmed the successful isolation of islet PROCR+ and duct cells, respectively (Figure 1E). The remaining acinar cells were mainly in the PROCR-, CD133- population, as shown by the high expression of the acinar marker *AMY1A* (Figure 1E). The PROCR+ population exhibited significantly higher expression of the signature genes of mouse Procr+ cells, including *RSPO1*, *FGF1*, *HOXA5*, *IGFBP5*, and *LGALS7* (Figure S1B), suggesting a conservation of the molecular signature of the PROCR+ population between species.

To establish a suitable culture medium for human PROCR+ cells, we initially employed enriched islet cells (post-Ricordi isolation and single-cell dissociation) to screen various small molecules and growth factors (Figure S3A-F). Through testing with numerous compound combinations based on the medium for mouse islet organoids^28^, we developed a 10-factor expansion medium capable of supporting islet-enriched cells to grow into colonies (see Figure S3G for its composition).

We then investigated the growth potential of various sorted subpopulations from enriched islet cells, i.e. PROCR+ (Live, Lin-, EpCAM+, PROCR+), PROCR- (Live, Lin-, EpCAM+, PROCR-) and duct cells (Live, Lin-, EpCAM+, CD133+) in this culture condition. Unsorted islet cells displayed a low colony-forming efficiency (16 out of 10,000 plated cells) (Figures S3B, S3E). In contrast, PROCR+ islet cells exhibited a 18-fold increase in clonogenicity (280 out of 10,000 plated cells), while PROCR-negative islet cells failed to form colonies (Figures 1F-H). CD133+ duct cells also readily formed colonies (520 out of 10,000 plated cells) (Figures 1F-H), highlighting the necessity of eliminating duct cells from such organoid culture. These observations were replicated across multiple donors (Figure 1H). Notably, organoids derived from endocrine or duct cells displayed distinct sizes and morphology. PROCR+ cell-derived organoids reached an average diameter of 110 μm by day 14 culture, exhibiting a dense spherical shape (Figures 1F-G). In contrast, CD133+ duct cell-derived organoids (ductal organoids) were significantly larger (approximately 180 μm diameter) and displayed a hollow, cystic morphology (Figures 1F-G), consistent with previous reports^38,60,61,63,64^. In PROCR+ cell-derived organoids, a portion of cells residing at the center of the organoid retained

PROCR expression (Figure 1I). In contrast, PROCR staining was completely absent in ductal organoids (Figure 1I). Both organoids expressed the pancreatic marker gene PDX1 (Figure 1J). Importantly, ductal marker SOX9 was absent in PROCR+ cell-derived organoids, whereas it was abundantly expressed in ductal organoids (Figures 1K). qPCR analysis of additional ductal markers, e.g., *PROM1*, *KRT7*, *CFTR*, *CD44*, and *HNF1B*, further demonstrated the differences in gene expression of these two organoids (Figures 1L). Interestingly, the expression of endocrine markers, e.g., *CHGA*, *CHGB* and *NEUROD1*, *PAX6* and *NKX2-2* were significantly higher in PROCR+ cell-derived organoids (Figure 1M), suggesting their endocrine differentiation potential.

We noted that PROCR staining was also observed in a small subset of acinar cells (Figure S1C). Therefore, we further analyzed acinar cells by FACS in post-Ricordi cell fraction with the acinar marker Ulex Europaeus Agglutinin-1 (UEA1) (Live, Lin-, EpCAM+, UEA1+)^65,66^ (Figure S1D-S1E), and determined that about 0.6% of UEA1+, PROCR+ cells existed in the acinar compartment (Figure S1F). Acinar cells are known to be refractory to growth in culture^67^. Nevertheless, to exclude the possibility that organoids originated from contaminating acinar cells, we sorted various acinar populations and verified that they could not form colonies under our culture conditions (Figures S1F–S1G). When UEA1+ cells were excluded from post-Ricordi fraction, the percentage of PROCR+ cells remained at approximately 5.6 ±0.8% of total EpCAM+ islet cells in FACS analysis (n=2 donors) (Figure S2B).

Collectively, we isolated PROCR+ cells from the pancreatic islet-enriched fraction and established an expansion medium to cultivate single PROCR+ islet cells into a distinct type of pancreatic organoid, which we refer to as PICO (PROCR+ cell-derived, Islet-resident, Clonogenic Organoids).

### Long-term propagation of functional human PICO *in vitro*

To assess the expandability of PICO, we implemented a passaging protocol. Organoids were maintained in the expansion medium and dissociated into single cells every 12-14 days for re-replating at a ratio of 1:2 to 1:10 under the same culture conditions (Figure 2A). This system maintained a stable average organoid size throughout serial passaging (Figures 2B), while enabling a 2- to 10-fold increase in organoid number with each passage (Figure 2C). PICOs could be passaged at least 10 times, spanning over 5 months. Starting from approximately 1,5000 organoids at passage 1, the number of organoids reached an average of 5x10^11^ organoids (∼2.5x10^14^ cells) by passage 10 (data from 7 donors), representing a 3x10^8^-fold expansion (Figure 2C). Notably, PICOs at the expansion stage can be cryopreserved after being dissociated into single cells, with an over 80% restoration rate upon thawing (Table S2).

**Figure 2.**
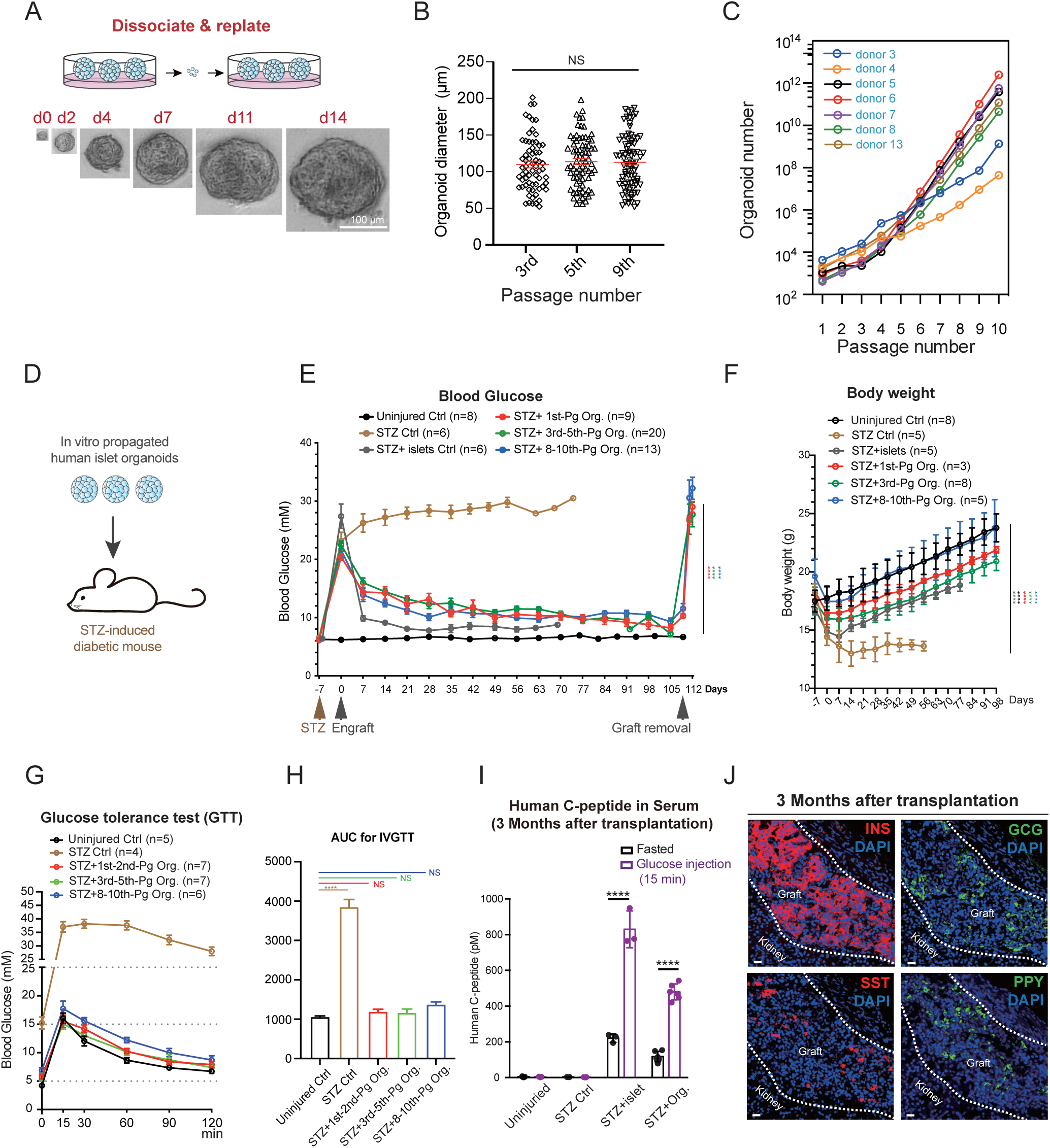
Long-term cultured islet organoids ameliorate mouse diabetes *in vivo*. (A) Illustration of organoid passaging strategy. Representative bright-field images of PROCR+ cell derived, Islet-resident, Colonogenic Organoids (PICO) formation (Donor #4). Scale bar, 100 µm. (B) Measurements of organoid sizes across different passages. Organoids sizes are shown as mean ± SEM. One-way ANOVA with Tukey’s test is used for the comparison of multiple groups. NS, p>0.05. Data were pooled from n=3 donors (see Table S1). (C) Cumulative yield of PICO from passage 1 (P1) to passage 10 (P10) derived from different donors (n = 7, color-coded, Table S1). Each symbol represents data derived from three technical replicates. (D) SCID-beige mice between 8-10 weeks old were induced diabetic with streptozotocin (STZ) injection 7 days before engrafting. Non-fasting blood glucose levels were monitored. (E) Human islet organoids from 1^st^-2^nd^ (early), 3^rd^-5^th^ (mid) or 8^th^-10^th^ (late) passages were engrafted under the kidney capsule of diabetic mice. Number of recipients for each group were indicated. Recipients non-fasting glucose levels were monitored weekly. Nephrectomy (graft removal) was performed at 3-month after transplantation, followed by glucose measurement. Data are presented as mean ± SEM. Two-way ANOVA with Dunnett’s test is used for comparison of multiple groups. **** p<0.0001. Data were pooled from n=8 donors (see Figure S4B, Table S1). (F) Body weight of mice before and after engrafting. Number of mice measured for each group were indicated. Data are presented as mean ± SEM. Two-way ANOVA with Dunnett’s test is used for comparison of multiple groups. Between groups, each group is compared with the STZ group. **** p<0.0001. Data were pooled from n=7 donors (see Table S1). (G) Glucose tolerance test was performed 1∼3 months after PICO engrafting. Recipients were fasted overnight, and glucose were injected intra-peritoneally. Numbers of mice analyzed per group are as indicated. Data are presented as mean ± SEM. Data were pooled from n=9 donors (see Table S1). (H) Respective AUC values of IVGTT results in (D). Data are presented as mean ± SEM. One-way ANOVA with Tukey’s test is used for comparison of multiple groups. Between groups, each group is compared with uninjured control group. NS p>0.05; **** p<0.0001. (I) Serum human C-peptide levels were measured by ELISA before or after (15 min after) glucose injection. n=3 mice for the normal, STZ ctrl or STZ+islet groups and n=6 mice for PICO engrafted group. Data are presented as mean ± SEM. Paired two-tailed t test is used for comparison. **** p<0.0001. Data were pooled from n=5 donors (see Table S1). (J) Representative immunostaining images of INS (β cells), GCG (α cells), SST (δ cells) and PPY (PP cells) on graft sites. Scale bar, 20 µm. Data were pooled from n=3 donors (see Table S1).

### Long-term cultured human islet organoids ameliorate mouse diabetes *in vivo*

To assess the in vivo functionality of PICO, they were engrafted into the kidney capsule of streptozotocin (STZ)-induced diabetic mice, which were subsequently monitored for glycemic improvement. A total of 800-1000 organoids (estimated to contain approximately 500,000 cells) from Expansion d14 culture at early (1^st^), mid (3^rd^-5^th^), or late (8^th^-10^th^) passage, or 300 freshly isolated human islets containing similar cell numbers (∼500,000 cells), were transplanted into each recipient mouse (Figure 2D). One-week post-transplantation, blood glucose levels were markedly reduced (Figure 2E), and body weight loss was reversed (Figure 2F). The glycemic improvement was maintained throughout the analysis period, with 62 mice analyzed at approximately 4 months (Figure 2E) and 11 mice analyzed at 1-year post-transplantation (Figure S4A). Notably, the organoid-transplanted groups, regardless of early, mid, or late passages, exhibited similar rescue effects as the fresh islet-transplanted group (Figure 2E). The rescue effects were consistent across the eight donor lines examined (Figure S4A-B). One-month post-transplantation, the mice transplanted with organoids exhibited significantly improved glucose clearance ability (Figure 2G-H). Human C-peptide was detected in the serum of organoid-recipient mice upon glucose challenge, reaching approximately 60% of the levels observed in fresh islet-recipient mice (Figure 2I). Removal of the transplants by nephrectomy resulted in an abrupt increase in blood glucose, confirming that the functional β cells resided in the kidney transplants (Figure 2E). Thus, the transplanted organoids were capable of secreting insulin and ameliorating hyperglycemia in a diabetic mouse model.

The grafted organoid cells were excised and analyzed after 3 months of transplantation. All four endocrine cell types were detected, including INS+ β cells, GCG+ α cells, SST+ δ cells, and PPY+ PP cells (Figure 2J). The host pancreases were also analyzed at 3 months post-transplantation and confirmed no signs of endogenous β cell recovery (Figure S4C-D). These observations further supported that the restoration of normal glycemia was due to the grafted PICO in the kidney.

### Differentiation of human PROCR+ organoids into functional endocrine cells

Next, we explored the endocrine-differentiation of PICO in culture. FACS analysis and immunostaining indicated that PICO at expansion day 14 expressed the pancreas marker PDX1 (Figure S5A), but lacked hormone expression (Figure 3B). Therefore, we developed a maturation protocol to facilitate their differentiation towards functional endocrine cells. Given the generally inverse relationship between cell proliferation and maturation during development^68,69^, we removed the growth-promoting factors CHIR99021 and SB202190. Notch signaling reduction, known to induce NEUROG3 expression and enhance endocrine cell formation^70–72^, has also been critical in generating β cells from ESCs^73,74^, and during acinar-to-β cell transdifferentiation^75^. Guided by these principles, we screened various media formulations (Figure S5B) and tested different treatment durations (day 7, day 12, and day 17). qPCR analyses revealed that media #8 at day 12 (12d) yielded optimal maturation, exhibiting the highest expression levels of endocrine lineage markers, *INS*, *GCG*, *NKX6.1*, *UCN3*, and *MAFA* among the tested conditions (Figure S5C). This formulation was designated ‘maturation medium’ (Figure S5B).

**Figure 3.**
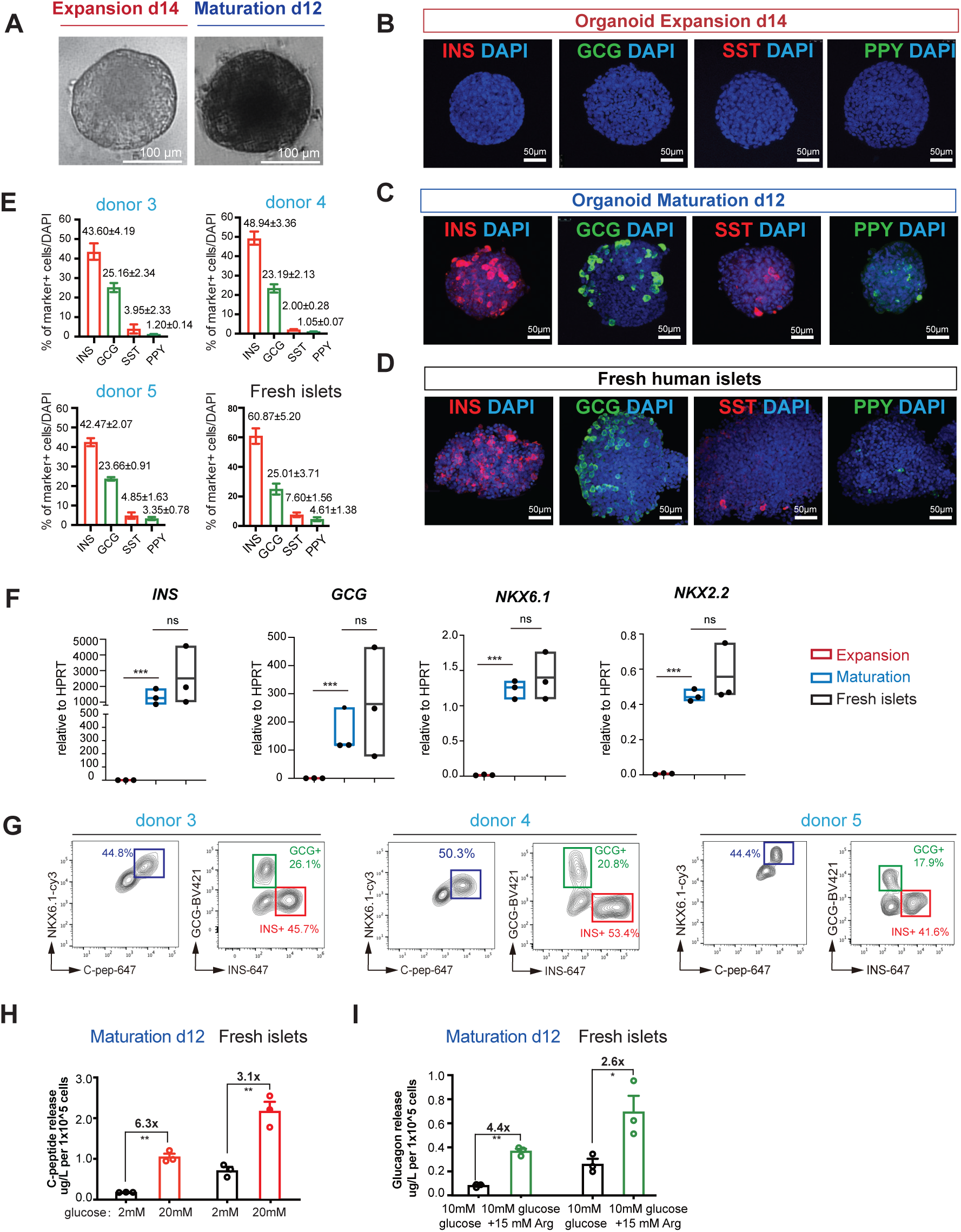
PICO exhibit full endocrine differentiation potential in vitro. (A) Representative bright-field images of PICO at expansion d14 or and at maturation d12 (Donor #4). Scale bar, 100 µm. (B-D) Whole-mount immunofluorescence analysis of pancreatic hormones in 3D organoids. Representative images (Donor #8) and quantification of INS+ (β cells), GCG+ (α cells), SST+ (δ cells), and PPY+ (PP cells) populations under expansion conditions (B), maturation conditions (C), and primary human islets (positive control; D; Donor #2). Scale bars, 20 µm. (E) Quantitative analysis of INS+ (β cells), GCG+ (α cells), SST+ (δ cells) and PPY+ (PP cells) cells in matured organoids, n=3 donors (Donor #3, #4 and #5). Data are presented as mean ± SEM from three technical replicates. The quantification data of fresh islets were combined from n = 4 donors (see Table S1). (F) RT-qPCR analysis of *INS*, *GCG*, *PDX1*, *NKX2.2* and *NKX6.1* mRNA expression levels in organoids under expansion condition and maturation condition, compared to primary human islets (control). Data were pooled from n=3 donors (see Table S1), presented as floating bars spanning minimal to maximum. Each symbol represents an organoid line, with data generated from three technical replicates. One-way ANOVA with Tukey’s test is used for the comparison of multiple groups. ns. p>0.05; *** p<0.001. (G) Flow cytometry analysis of β-cell maturation markers (C-peptide+/NKX6.1+) and endocrine lineage commitment (INS+ or GCG+) in matured organoids from different donors (n=3). Data represented as average of three technical replicates. (H) ELISA measurements of human C-peptide secretion from matured PICO (left) and primary human islets (right) following sequential stimulations with low (2 mM) or high (20 mM) glucose. Data were combined from n=3 donors (see Table S1). Each symbol denotes data from one donor, averaged over three technical replicates. Data are presented as mean ± SEM. Paired two-tailed t test is used for comparison. ** p<0.01. (I) ELISA measurements of human glucagon secretion from matured PICO (left) and primary human islets (right) following sequential stimulations with 10 mM glucose (basal) or 10 mM glucose plus 15 mM arginine (Arg). Data were combined from n=3 donors (see Table S1). Each symbol denotes data from one donor, averaged over three technical replicates. Data are presented as mean ± SEM. Paired two-tailed t test is used for comparison. * p<0.05; ** p<0.01.

While organoid size progressively increased during expansion, it remained stable during maturation (Figure 3A). Intriguingly, the organoids appeared darker post-maturation (Figure 3A, Figure S5D). Notably, transitioning PICO from expansion to maturation conditions resulted in an approximate 60% loss of organoids at maturation-d12, as reflected by reduced organoid density and total cell counts (Figure S5D-E). This loss coincides with the withdrawal of key growth-promoting factors—including CHIR and SB202190—which are essential for expansion but are removed to promote endocrine differentiation, and the physical manipulation required to release and replate the organoids in fresh matrix (see Methods). The observation that the size of the remaining organoids remained stable suggests that, once established, these surviving organoids retain their structural integrity under maturation conditions.

Immunostaining of organoids at maturation-d12 revealed that approximately 43.6% ± 4.2% of cells expressed insulin, 25.2% ± 2.3% expressed glucagon, and smaller proportions expressed somatostatin (SST, 4.0% ± 2.3%) or pancreatic polypeptide (PPY, 1.2% ± 0.1%) (Figure 3C, 3E). Notably, ∼25% cells remained hormone-negative. This endocrine composition, consistently observed across multiple donors (Figures 3E), fell within the range of proportions seen in primary human islets of similar size (Figure 3D, 3E). Further assessment of organoid maturation by qPCR analysis of endocrine markers demonstrated that the expression levels of *INS*, *GCG*, *PDX1*, *NKX2.2*, and *NKX6.1* in PICO at maturation-d12 were comparable to those in fresh human islets and, as expected, significantly higher than in PICO at the expansion stage (Figure 3F). To quantify the percentage of β cells, we assessed the co-expression of NKX6.1 and C-peptide, revealing β cell percentages ranging from 41% to 53% (Figures 3G). Further analysis of INS and GCG expression corroborated the β cell proportions observed in immunostaining and indicated that the percentage of α cells ranged from 17% to 26%. Importantly, no cells co-expressing INS and GCG were detected (Figures 3G). These findings were replicated across multiple donors (Figures 3G).

Moreover, PICO at maturation-d12 exhibited rapid increases in cytosolic Ca2+ concentrations in response to glucose, followed by a return to baseline (Figure S5F), indicating their ability to respond acutely to glucose stimulation. Transmission electron microscopy (TEM) confirmed the presence of insulin granules in maturation-d12 organoids (Figure S5G). ELISAs revealed glucose-stimulated human C-peptide secretion at ∼50% of fresh islet capacity (Figure 3H), alongside arginine-stimulated glucagon release (Figure 3I). Together, these results demonstrate that our maturation protocol drives PICO differentiation into functionally competent endocrine organoids, recapitulating hallmark features of mature islets, including hormone production, dynamic glucose responsiveness, and regulated insulin/glucagon secretion.

### Single-cell RNAseq analysis reveals full differentiation potential and a progenitor-like population in PICO

To determine the molecular characteristics of PICO, we conducted single-cell RNA sequencing (scRNA-seq) on organoid cells at Expansion d14 and Maturation d12 (Figures 4A and S6A). We integrated these data with primary human islet cell datasets, including six published datasets^76–81^ and two newly generated datasets (Figures 4A and S6B). During Expansion, two distinct organoid populations (Org.1-2) were identified (Figure 4A). At Maturation, in addition to the existing organoid populations, α-, β-, δ-, and PP-like cells emerged, clustering with primary α, β, δ, and PP cells, respectively (Figure 4A and S6C). To assess PICO cell maturation, we compared the expression of hormone genes (INS), islet cell markers (IAPP), and transcriptional regulators (PDX1, NKX6.1, MAFA) between PICO-β cells and primary β cells, finding comparable expression levels (Figure 4C). Similarly, marker and hormone gene expression levels in PICO -α cells, PICO -δ, and PICO -PP cells were comparable to those in primary cells (Figure 4C). It is noteworthy that immunostaining (IF) and FACS-methods widely used in the field identify INS+ cells based on hormone expression or marker positivity, estimating ∼50% INS+ cells in PICO (Figure 3C, 3E, 3G). In contrast, scRNA-seq criteria define β-cell identity through the co-expression of thousands of genes characteristic of primary human β-cells, a far more stringent standard. By this measure, ∼24% of PICO-β cells transcriptionally overlap with primary β-cells (Figure 4A). This discrepancy arises from differing definitions of maturity: IF and FACS confirm functional maturity (hormone production), while scRNA-seq assesses molecular maturity (global transcriptional fidelity to native β-cells).

**Figure 4.**
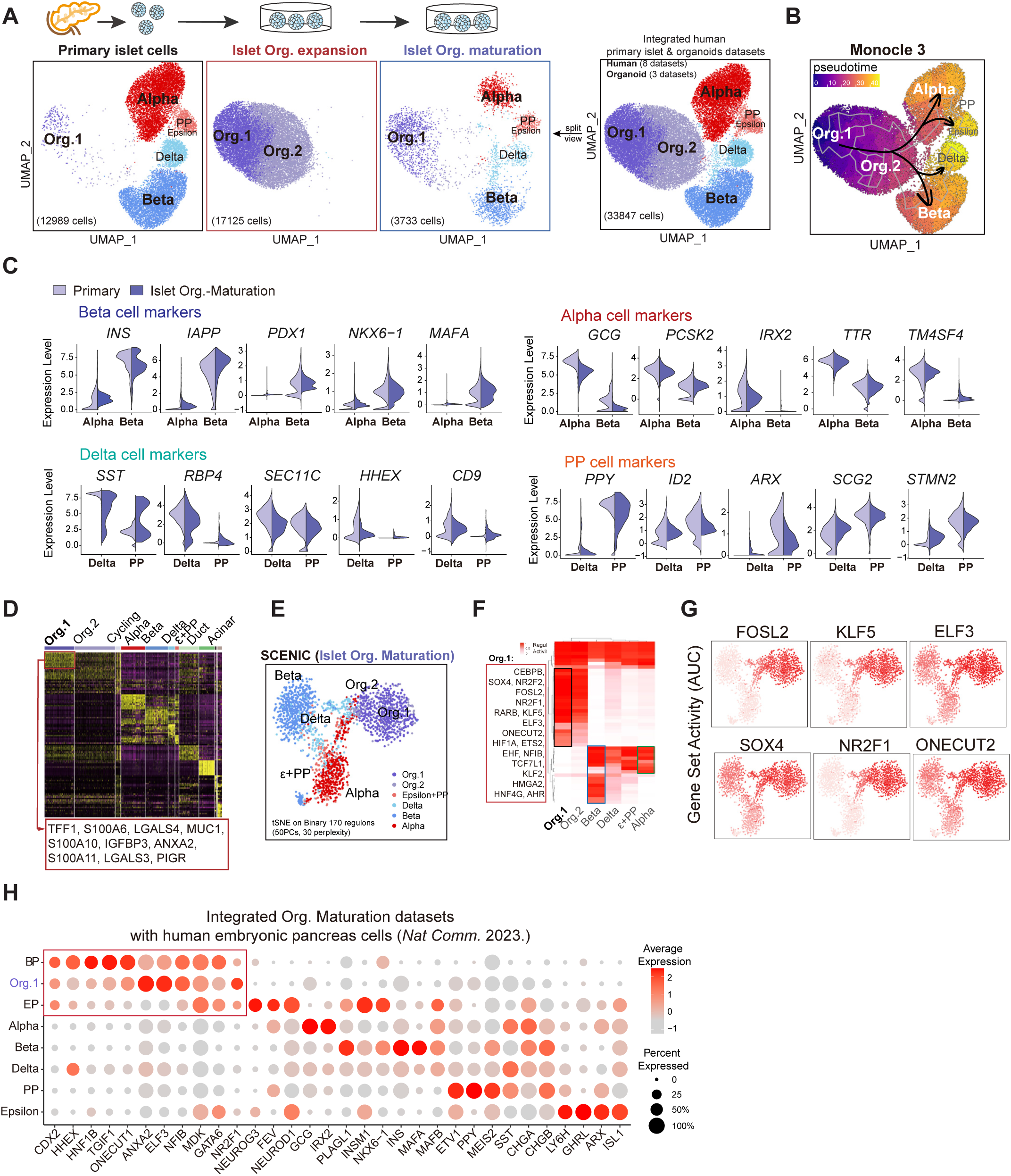
Single-cell RNAseq analysis reveals full differentiation potential and a progenitor-like population in PICO. (A) Cultured organoids in expansion (donor #3) and maturation (donor #13) single-cell (sc) RNA-seq dataset were integrated with primary human pancreatic islets (donor #1 and #2) scRNA-seq datasets and projected onto UMAP plots, colored by cluster assignment, and annotated post hoc. Both the aligned (right) and split (left) views are shown. Cell counts of each dataset are labeled in brackets. (B) Developmental trajectory produced by Monocle 3. Pseudotime (arbitrary units) is depicted in purple to yellow in brackets. (C) Violin plots comparing the expression levels (log_2_(TPM+1)) of representative genes of α, β, δ and PP cell population in matured PICO and human pancreatic islets. (D) Heatmaps of different cluster-enriched genes. Each column represents a single cell and each row represents one signature gene. The colors ranging from purple to yellow indicate low to high relative gene expression levels. (E) SCENIC t-SNE plots illustrated mature organoid cells, color-coded by cell types, using the binary regulon activity matrix. (F) The heatmap depicted the scaled activity of the top-scoring regulons across cell types, derived from the same activity matrix. (G) SCENIC t-SNE plots showing several active regulons in Org.1 population, cells were colored according to the corresponding binary regulon activity. (H) Cultured organoids (in maturation) scRNA-seq dataset are integrated with human embryonic pancreas scRNA-seq datasets (endocrine cells, EPs and and BPs at PCW 7-11). Dotplot for signature genes of each cluster. The shadings denote average expression levels, and the sizes of dots denote fractional expression.

Monocle3 analysis revealed a differentiation trajectory from Org.1 to Org.2 and eventually to α-, PP- and β-, δ-like cells, suggesting that PICO cells originate from Org.1 (Figures 4B and S6D). Org.2 cells might represent a transitional population. The near absence of Org.2 cells at later time points is consistent with their role as a transient intermediate that gives rise to hormone-expressing cells (Figures 4A, 4B).

Interestingly, a population of Org.1-like cells was found in primary islet datasets (Figure 4A), suggesting that this progenitor-like state represents a bona fide islet-resident population rather than an in vitro artifact. Heatmap analysis of differentially expressed genes (DEGs) highlighted the signature genes of Org.1 (Figure 4D). To define Org.1’s gene regulatory networks, SCENIC analysis identified activated transcription factors (TFs) (Figure 4E-G, S6E). The activation of FOSL2, KLF5 and ELF3 may indicate Org.1’s proliferative state, while SOX4, NR2F1, and ONECUT2 activities suggest a similarity to both fetal bipotent progenitors (BPs, also called trunk progenitors) and endocrine progenitors (EPs) ^82,83^(Figure 4G). When aligning PICO data with published datasets of developing human pancreatic endocrine cells, including EPs and BPs at fetal week 7-11^82^, we observed significant transcriptional overlap among BPs, Org.1 and EPs. Transcription factors including CDX2, MDK, GATA6, and NR2F1 were expressed at high levels in all three groups, underscoring their molecular resemblance (Figure 4H).

Surprisingly, PROCR levels appeared low in abundance in our scRNA-seq analysis, likely due to the inefficiency of the PROCR primer on the 10X platform (Figure S7A). Nonetheless, PROCR expression in PICO was clearly detected by immunostaining (Figure 1I), and confirmed by FACS analysis and qPCR validation (Figure S7B-D). When PROCR+ and PROCR- cells were FACS-isolated and replated, only PROCR+ cells could form colonies in serial passage, supporting that PROCR+ cells are the origin of the organoids (Figure S7E). qPCR analysis further demonstrated that PROCR+ cells isolated from PICOs show high expression of Org.1 signature genes (Figure S7F), indicating their resemblance to Org.1 cells.

Collectively, the PICO system provides a unique opportunity to identify a progenitor-like population (Org.1) in the adult human pancreas, with molecular features reminiscent of both fetal BPs and EPs. Importantly, these cells can efficiently generate mature endocrine cells in vitro.

### The progenitor-like population shares ductal features

A population of Org.1-like cells was identified in primary islet datasets through PICO data integration (Figure 4A). Interestingly, excluding PICO data from the integration analysis resulted in the annotated Org.1-like cells clustering with duct cells in the human primary pancreas dataset (Figure S8A-A’), suggesting molecular similarities between Org.1 cells and duct cells. However, Gene Ontology (GO) analysis of DEGs revealed distinct differences: Org.1 cells are preferentially enriched for genes involved in fatty acid, prostaglandin, and retinol metabolism process, as well as beta cell and epithelial cell proliferation (Figures S8B). Violin plots of Org.1 signature genes further illustrated their distinction from duct cells (Figure S8C). Analysis of the ductal organoid dataset ^84^ demonstrated that PICO and ductal organoids are completely different (Figures S8F-G).

Among the signature genes of Org.1 (Figure 4D), we noted that *TFF1* and *S100A6*, have been previously described in an “activated/migrating progenitor-like” ductal cell subset^85^. Indeed, PICO cells exhibit some ductal cell features but have much lower expression levels of duct markers (*SOX9*, *DCDC2*, *HES4*, and *CFTR*), which are absent in endocrine cells (Figure S8D). Importantly, they also showed low expression of endocrine markers (*CHGA*, *CHGB*, *TM4SF4* and *NEUROD1*) (Figure S8E), in line with qPCR analysis shown in Figure 1M. These observations raise the possibility that Org.1 cells might have a developmental origin from fetal truck, with a differentiation propensity towards the endocrine lineage.

### Long-term cultured human islet organoids ameliorate non-human primate diabetes in vivo

As a proof-of-principle for clinical translation, we evaluated PICO in a non-human primate (NHP) model of diabetes (Figure 5A). Two healthy *Cynomolgus macaques* were rendered diabetes by a single-dose STZ injection (Figure 5B, 5C). Exogenous insulin administration was initiated two days later, and fasting glucose levels (red line) along with daily insulin usage (blue line) were recorded (Figure 5B, 5C). Both animals achieved a stable diabetic state within five weeks, characterized by fasting blood glucose levels exceeding 20 mM and daily insulin requirements of 6-10 IU, consistent with published NHP models^86–88^. Impaired glucose clearance was confirmed by intravenous glucose tolerance test (IVGTT) (Figure 5D, 5E), and HbA1c levels progressively increased to 10.1% (recipient #1) and to 8.1% (recipient #2) by nine weeks post-STZ induction (Figure 5J, 5K).

**Figure 5.**
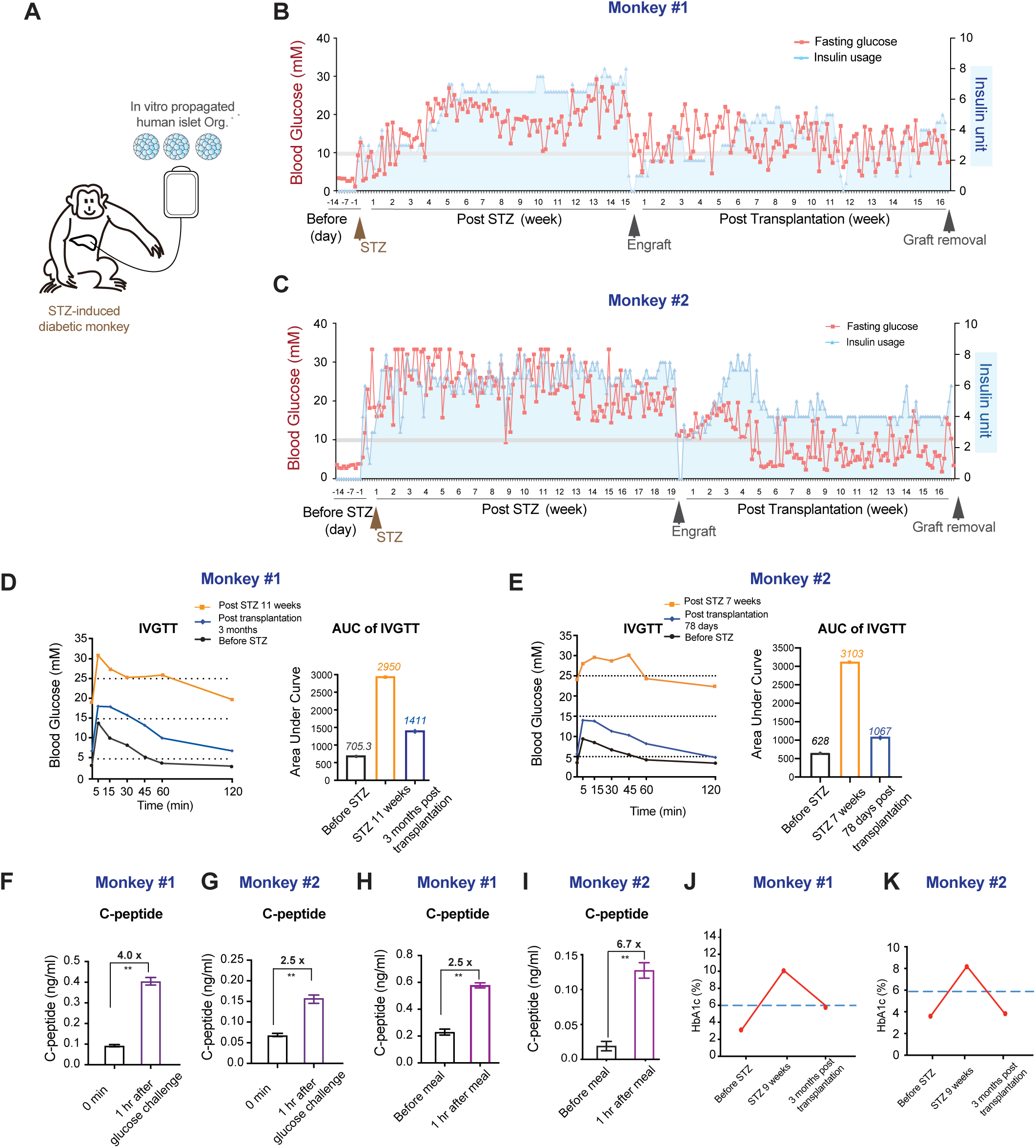
PICO ameliorate diabetes in non-human primate. (A) Illustration on human PICO transplantation into STZ-induced diabetic *cynomolgus* macaque model. Organoids used for monkey transplantation was derived from Donor #11 (Passage 4) and Donor #15 (Passage 6) (see Table S1). (B, C) Daily fasting blood glucose levels (red line) and daily exogenous insulin usage (blue shielding) in two macaques before and after infusion of PICO (transplantation procedure conducted at week 0). (D-E) Blood glucose levels and respective AUC values in response to glucose challenge (IVGTT) in two macaques: before STZ induction (black line); after diabetic induction (11 weeks after STZ induction, orange line); and after PICO transplantation (3 months for D and 78 days for E after infusion, respectively, navy blue line). (F-I) C-peptide secretion of two macaques 1 hour after glucose challenge (F, G) and 1 hour after meal intake (H, I). Data are presented as mean ± SEM of three technical replicates. Unpaired two-tailed t test is used for comparison. ** p<0.01. (J, K) HbA1c levels of two macaques before STZ induction, 9 weeks after diabetic induction and 3 months after PICO infusion. Blue dotted line indicates normal HbA1c range.

To permit xenotransplantation, both animals received immunosuppressive therapy (tacrolimus and sirolimus) beginning nine days prior to surgery (Figure S9A-E). On the day of transplantation, each macaque received a single intraportal infusion of 15,000 islet equivalents (IEQ)/kg of PICO. Fasting glucose level and insulin usage were recorded daily for 16 weeks (Figure 5B, 5C). Both recipients exhibited significantly reduced fasting blood glucose and a 50% reduction in exogenous insulin requirements, averaging 4 IU (Figure 5B, 5C). Within three months post-transplantation, IVGTT demonstrated markedly improved glucose clearance in both animals (Figure 5D, 5E). This was accompanied by increased stimulated C-peptide secretion following glucose challenge (4-fold in recipient #1 and 2.5-fold in recipient #2) (Figure 5F, 5G), as well as enhanced post-meal C-peptide responses (2.5-fold in recipient #1 and 6.7-fold in recipient #2) (Figure 5H, 5I). Importantly, in recipient #1, HbA1c decreased from 10.1% pre-transplantation to 5.9% three months post-transplantation (Figure 5J); in recipient #2, HbA1c declined from 8.1% to 3.6% over the same period (Figure 5K). Throughout the experiment, both recipients behaved normally without severe adverse responses, and body weight remained within the normal range (Figure S9F-G). Liver function was monitored frequently, showing normal levels of alanine aminotransferase (ALT), aspartate aminotransferase (AST), and total bilirubin (Figure S9H-M). Both macaques were euthanized 16 weeks post-transplantation, and a full autopsy showed no evidence of graft outgrowth or cell migration into other organ sites (Figure S9N-Q).

Collectively, our data from two independent diabetic macaques demonstrate that PICO transplantation improves insulin secretion and achieves sustained glycemic control, representing a critical milestone toward clinical translation.

## Discussion

In this study, we identify PROCR+ cells within human adult pancreatic islets and establish a defined culture system to generate functional, self-organizing endocrine organoids (PICO). PICO at maturation stage recapitulate the multicellular architecture of native islets, containing glucose-responsive α-, β-, δ-, and PP cells in physiological ratios, and exhibit dynamic insulin/glucagon secretion. Remarkably, PICO achieve long-term exponential expansion in vitro and rapidly ameliorate hyperglycemia in diabetic mice and macaque post-transplantation, positioning them as a scalable therapeutic cell source.

The existence of stem/progenitor cells in the adult pancreas has been debated for decades. Earlier pioneering studies implicated ductal cells as facultative progenitors, with evidence suggesting that ductal cells are capable of endocrine differentiation under specific conditions ^25,89,90,44,62,64^. However, murine lineage tracing later emphasized β cell replication as the primary renewal mechanism^26^. While these findings did not exclude facultative progenitors, they ignited a debate about the relevance of adult pancreatic stem cells in homeostasis and regeneration. Recent work adds to the body of research by identifying multipotent progenitors in both ducts^42,91–93^ and islets^27,28^, and proposing context-dependent regeneration: β-cell replication maintains homeostasis, while metabolic stress or injury activates dormant progenitors^25^.

Our discovery of PROCR+ islet progenitors provides new insights that help contextualize this debate. Although PROCR+ cells share certain molecular features with ductal cells, PROCR+ cells are transcriptionally primed for endocrine differentiation. In contrast to human duct-derived organoids, which typically retain exocrine/ductal programs and exhibit limited endocrine potential^38–44^, PICO derived from PROCR+ cells self-organize into islet-like structures containing multiple endocrine cell types. Single-cell transcriptomics further revealed that PROCR+ cells constitute a progenitor population (Org. 1) that may have previously been misclassified as ductal cells, potentially explaining why this population has remained undetected without specific markers and functional assays. The concept of quiescent progenitor populations that remain largely inactive under homeostatic conditions is well established in other slowly renewing tissues, such as muscle^94^ and neural systems^95^, and it is plausible that islet-resident progenitors may behave similarly.

We acknowledge that despite our efforts to exclude acinar and ductal cells using UEA-1 (acinar marker) and CD133 (ductal marker) depletion, and with the sorting purity of PROCR+ cells routinely exceeds 95%, we cannot entirely rule out the possibility of rare contaminating cells. Nevertheless, our data consistently show that PROCR+ cells are enriched within the islet fraction relative to exocrine tissue. This enrichment, together with their intra-islet localization by immunofluorescence, supports the presence of a bona fide PROCR+ population within human islets. This does not preclude the existence of rare PROCR+ cells in exocrine compartment, and future studies aimed at identifying such a population are warranted.

While significant progress has been made in the directed differentiation of ESCs/iPSCs into β-like cells^73,74,96–108^, these systems continue to face challenges, including tumorigenic risks from residual pluripotent cells, variable differentiation efficiency, and the presence of off-target lineages, such as enterochromaffin-like cells, which can affect functional purity^109–112^. Adult stem cells (ASCs) present a safer alternative due to their reduced tumorigenicity and ethical acceptability^113,114^, though their limited expansion capacity has historically restricted their clinical scalability. Our work presents an alternative paradigm by leveraging PROCR+ progenitors-derived PICO, which uniquely combine the safety of ASCs with unprecedented *in vitro* expandability. This platform offers new possibilities for the scalable generation of human islet tissue.

Transplantation of PICO into diabetic mice and non-human primates ameliorates hyperglycemia within two weeks, halting disease progression. This rapid efficacy likely reflects the adult epigenetic state of PICO, which bypasses the prolonged maturation required by ESC/iPSC-derived cells. However, PICO’s *in vivo* functional maturation lagged behind primary islet transplants, highlighting opportunities to refine pre-transplant protocols. To our knowledge, this work represents the first demonstration of ASC-derived islet organoids ameliorating diabetes in primate models—a critical step toward clinical translation. Future studies will optimize *in vitro* maturation and define molecular checkpoints governing post-transplant integration.

In summary, this study identifies PROCR+ cells as a reservoir of endocrine progenitor potential within adult human islets and establishes PICO as a scalable, physiologically relevant platform for diabetes research and therapy. By recapitulating native islet architecture and function, PICO advances drug discovery, disease modeling, and regenerative strategies, positioning adult progenitor-derived organoids as a transformative tool in pancreatic biology.

## Limitations of the study

While this work establishes PICO as a promising therapeutic platform, two key limitations warrant further investigation. First, current *in vitro* culture conditions incompletely mimic the native pancreatic niche, resulting in a 60% reduction in organoid density during the 12-day maturation phase. Notably, surviving organoids retain stable size, suggesting attrition occurs at the whole-organoid level rather than via individual cell loss. Single-cell transcriptomics further identified subpopulations of cells with immature transcriptional profiles at harvest, indicating incomplete maturation under existing protocols. Second, the reliance on Matrigel presents practical challenges: its high cost and labor-intensive handling make it impractical for large-scale organoid production, while regulatory concerns regarding its non-human components complicate clinical-grade translation. Future studies should prioritize staged nutrient supplementation to enhance survival and explore synthetic or gel-free matrices for clinical-grade organoid production.

PICO’s efficacy in reversing diabetes in a preclinical primate model underscores its therapeutic potential and provides a critical foundation for advancing this technology toward clinical trials. By addressing attrition and scalability, this platform holds transformative promise for redefining regenerative strategies for diabetes.

## ACKNOWLEDGMENTS

Thanks to Dr. Chi Chung Hui of University of Toronto for helpful comments on the manuscript. This research was supported by grants from the National Key Research and Development Program of China (2020YFA0509002 to Y.A.Z.), the National Natural Science Foundation (32588301 to Y.A.Z.), the CAS Project for Young Scientists in Basic Research (YSBR 014 to Y.A.Z.), Science and Technology Commission of Shanghai Municipality (23J21901400 to Y.A.Z.), Shanghai Municipal Science and Technology Major Project, CAS Croucher Funding Scheme for Joint Laboratories (CAS24801 to Y.A.Z.; CAS24SC02 to T.H.C. and Y.A.Z), and New Cornerstone Science Foundation (NCI202254 to Y.A.Z.).

## AUTHOR CONTRIBUTIONS

Y.A.Z. designed the project and wrote the manuscript. D.W., Q.C.Y. and W.S. developed the expansion protocol. W.S. developed the maturation protocol. W.S. and C.L. performed single-cell sequencing and bioinformatic analysis. W.S. and D.W. performed immunostaining, TEM, and ELISA. W.S., J.C. and D.Z. performed FACS analysis. D.W., Q.C.Y. and Y.X performed mouse transplantation and diabetic phenotype analysis. S.W., B.Z., Q.C.Y., W.S. and Y.S., performed NHP transplantation and diabetic phenotype analysis. S.S., H.B., W.W., L.L. A.S., Y.C., W.G., J.W., and H.F. obtained ethical approval for the use of human tissue and collected and processed the human samples. Q.C.Y., S.S. and L.L. helped with manuscript editing.

## STAR⍰Methods

### Key resource table

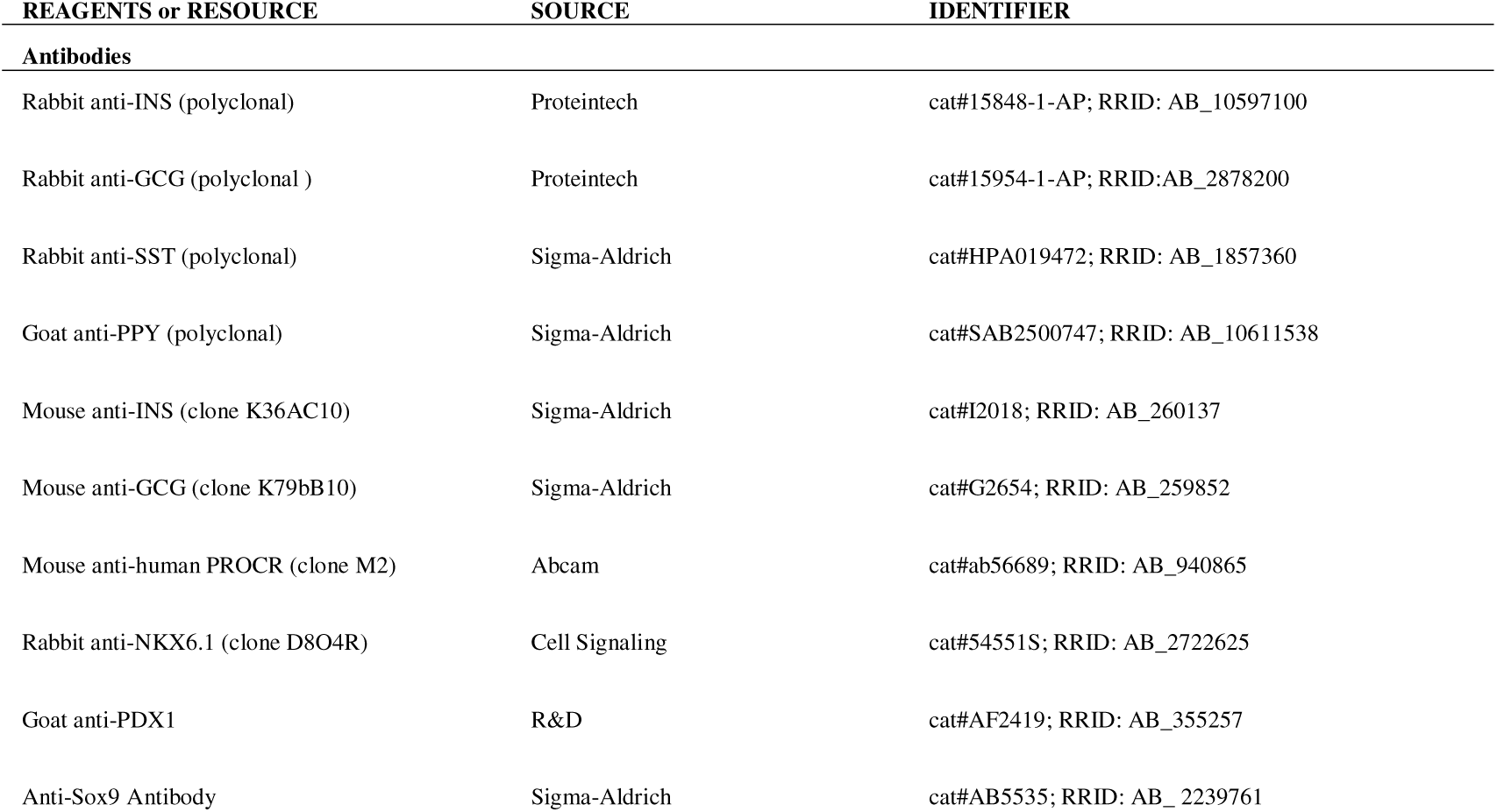
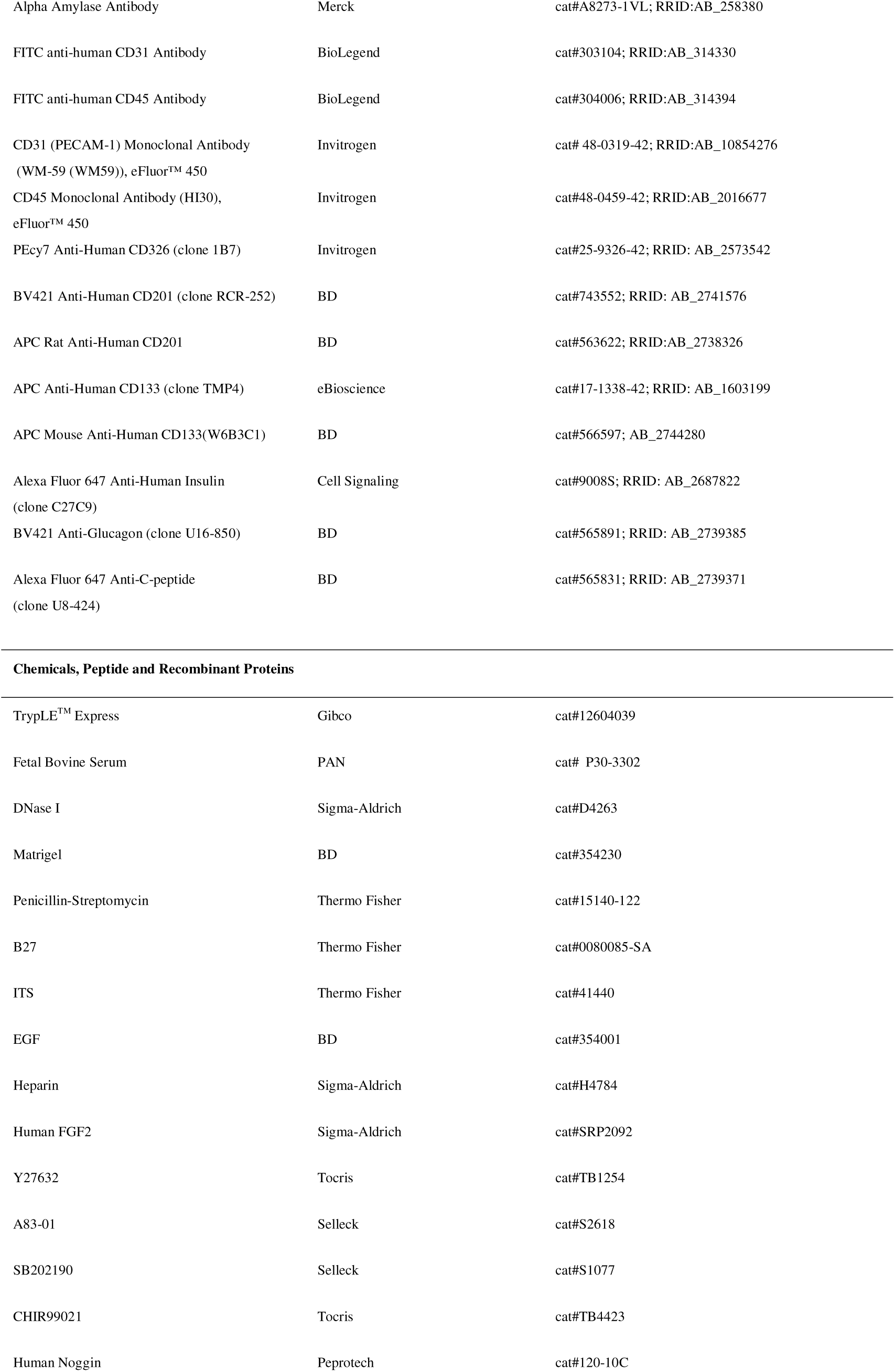
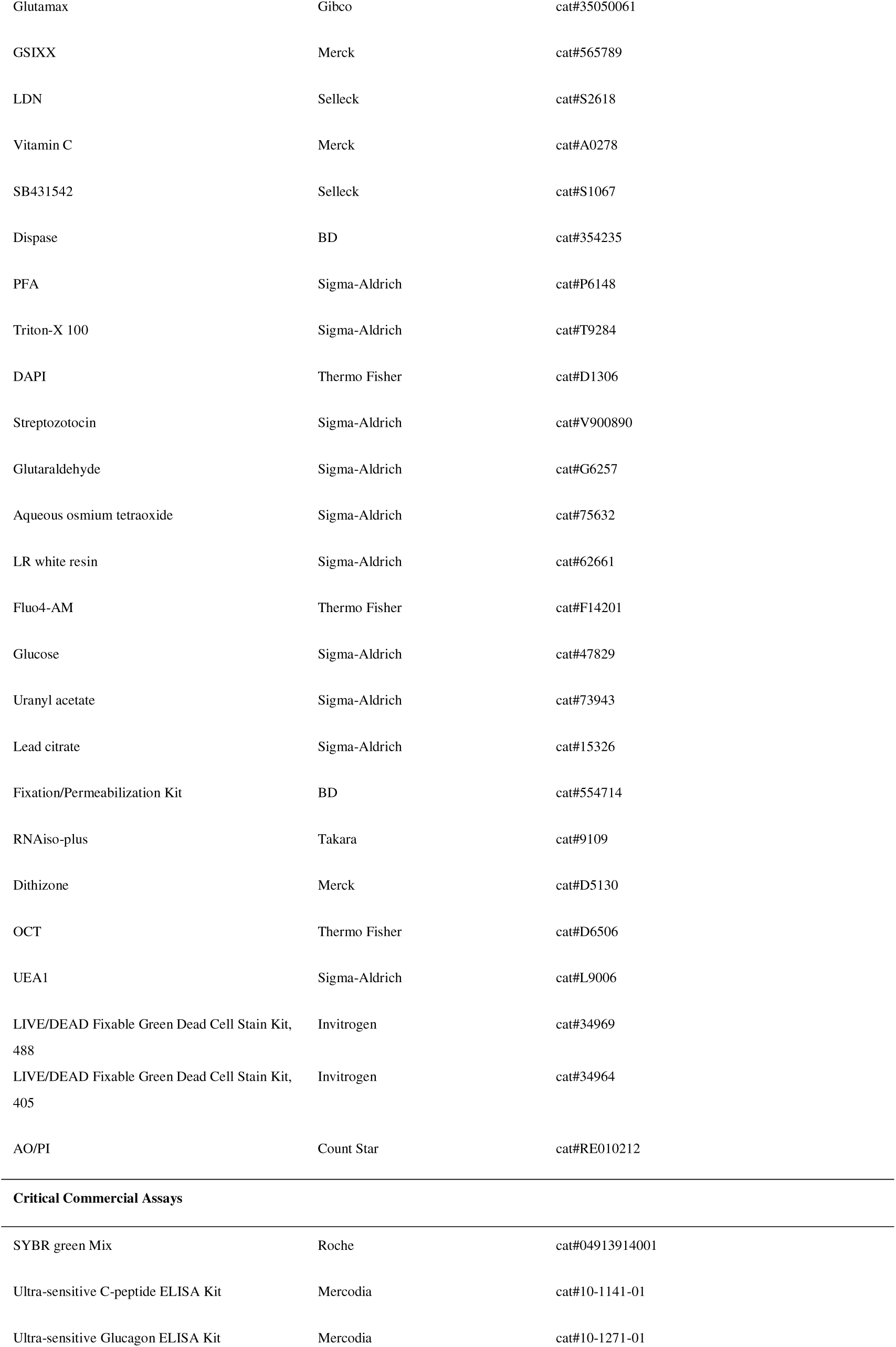
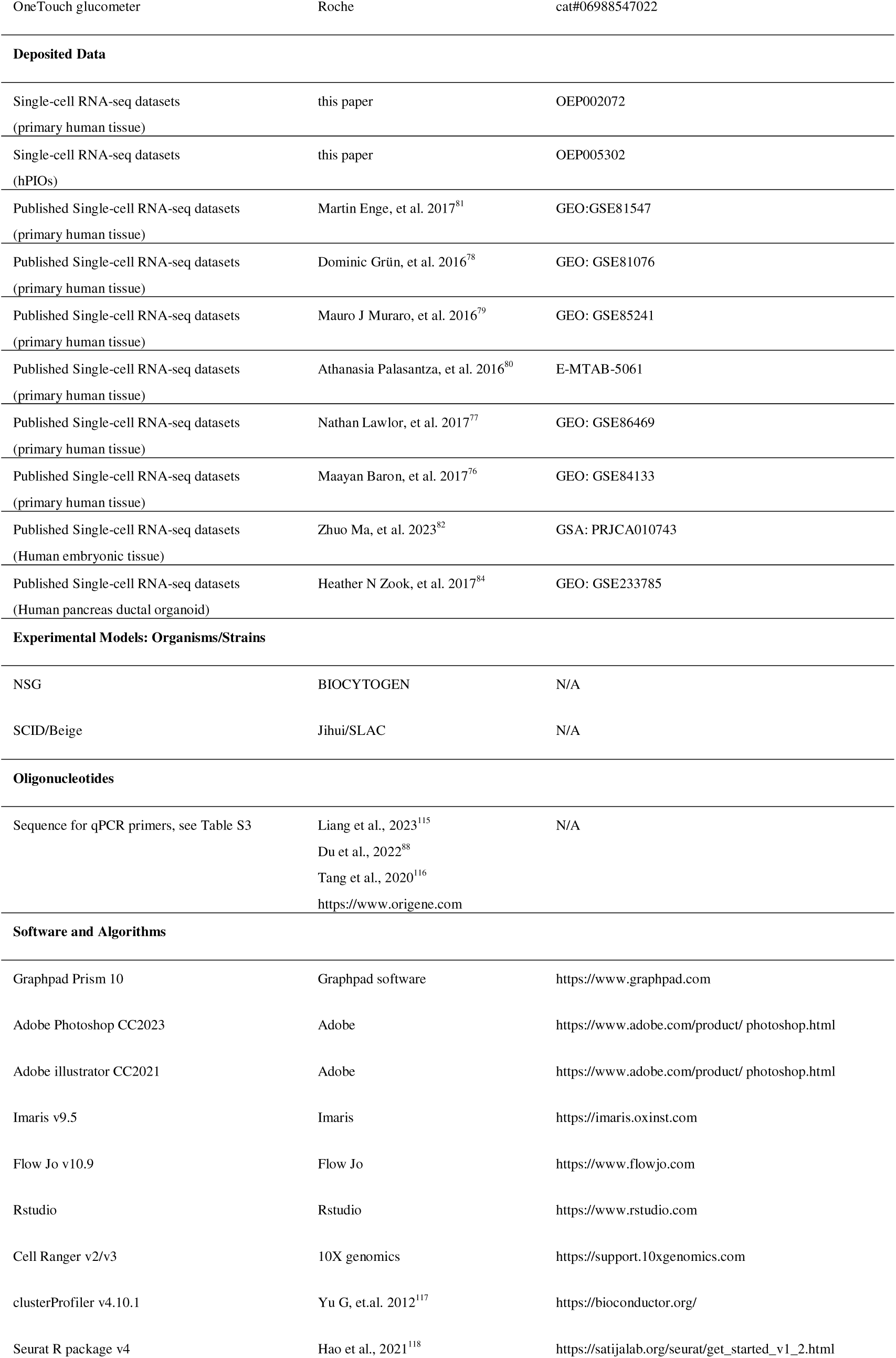
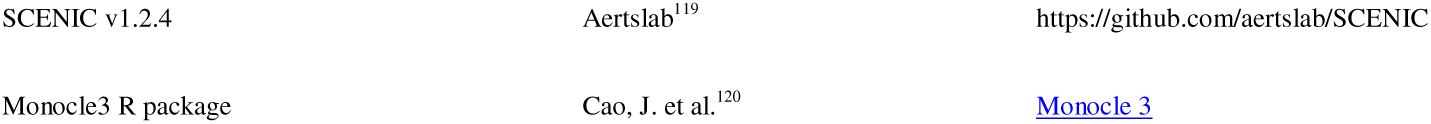

## Resource availability

### Lead contact

Further information and requests for resources and reagents should be directed to and will be fulfilled by the Lead Contact, Yi Arial Zeng.

### Materials availability

All unique/stable reagents generated in this study are available from the Lead Contact with a completed Materials Transfer Agreement.

### Experimental model and subject details Mice

NSG or SCID/Beige mice were used for organoids/islets transplantation study in this paper. All the animal experimental procedures were approved by the Animal Care and Use Committee of Center for Excellence in Molecular Cell Science, Chinese Academy of Sciences, with a project license of IBCB0065.

### Patients and clinical specimens

All human pancreas specimens were obtained from Ruijin Hospital, Shanghai Jiaotong University School of Medicine (Ethics approval number: 2022375), and Zhongshan Hospital, Fudan University School of Medicine (Ethics approval number: B2023-037R). The use of samples for research was approved by ethical committees and informed consent was obtained from donors.

## Method details

### Islets enrichment and purification

Human islet enrichments were performed at the collaborating hospitals with standard protocols (whole pancreas perfusion followed by density gradient centrifugation). The purity of the islets was assessed by DTZ staining. The purities range from 30-80%.

### Preparation of human pancreas single cell suspension and FACS sort

Human pancreatic islets were dissociated into single cells by enzymatic digestion. Briefly, islet enrichments were incubated with 2-4 ml TrypLE^TM^ Express enzyme and incubated at 37°C in a humidified 5% CO_2_ incubator for 6 min, followed by addition of 0.1 mg/ml DNase I and further incubation for 3 minutes. During the process, gentle manual pipetting was performed to disrupt aggregates. The enzymatic reaction was terminated by adding 10 ml PBS supplemented with 5% FBS. Cells were then filtered through a 70 μm cell strainer. The resulting single-cell suspension was washed twice with PBS and resuspended in appropriate volume of buffer.

For FACS analysis and sorting, cells were stained with LIVE/DEAD Fixable Dead Cell Stain Kit and dead cells were gated out in all analysis. Antibody incubation was performed on ice for 30 minutes at a dilution of 1:200. After two washes, cells were filtered through a 40-μm cell strainer. Sorting experiments were performed using FCAS Melody or FACS JAZZ machine (Becton Dickinson).

For UEA1+ cell labelling and isolation, human islet enrichments were resuspended in serum-free CMRL 1066 medium, and incubated with FITC-conjugated Ulex Europaeus Agglutinin 1 (UEA1) at a final concentration of 100 ug/ml for 1hour at 37□ in a humidified 5% CO_2_ incubator. Following incubation, cells were washed twice with PBS to remove unbound lectin, dissociated into single cells, followed by incubation with antibodies and cell sorting.

The purity of the sorted population was routinely checked and ensured to be more than 95%.

### Culture human islet organoids

A 48-well plate was coated with 100 μl of Matrigel per well and incubated at 37°C in a humidified 5% CO_2_ incubator until solidified. FACS sorted PROCR+ human pancreatic cells were resuspended in 500 μl/well of expansion medium (IMDM/F12 supplemented with1% Penicillin-Streptomycin, 2% B27, 1% ITS, 50 ng ml^-1^ EGF, 2 μg ml^-1^ heparin, 10 ng ml^-1^ FGF2, 2.5 μM CHIR99021, 0.5 μM A83-01, 100 ng ml^-1^ Noggin, 2.5 μM Y27632, 0.5 μM SB202190) and plated onto the Matrigel-coated wells. The medium was refreshed every 2-3 days.

The organoids were passaged every12-14 days. For passage, organoids were released from the Matrigel by incubation with dispase at 37°C for 30 minutes. The organoids were dissociate into single cells by digestion with TrypLE^TM^ Express at 37°C for 5-8 minutes. The single cells or cell clusters were then resuspended and replated onto Matrigel coated plates at a split ratio of 1:2-1:10.

For maturation, the organoids were released from Matrigel by incubation with dispase at 37°C for 30 minutes. The organoids were resuspended and replated onto Matrigel coated plates, supplemented with maturation medium (CMRL1066 supplemented with 1% Penicillin-Streptomycin, 2% B27, 1% ITS, 0.5 μM A83-01, 2.5 μM Y27632, 100 nM GSIXX and 0.25 μM LDN193189). The medium was refreshed every 3 days.

### Cryopreservation and recovery of PICO

PICO were dispersed using TrypLE^TM^ Express. Released single cells were rinsed with 5% FBS and spun down at 250g for 5 min. The cells were cryopreserved at a concentration of 1-5x10^6 cells per ml in cryopreservation medium and aliquoted into 2 ml vials, with 1 ml per vial. The vials were initially stored at -80°C for 24 hours before being transferred to liquid nitrogen for long-term storage.

Cryopreserved vials were thawed in a 37□ water bath. Cell suspension was then transferred into a 15 ml centrifuge tube containing 10 ml medium and centrifuged at 250g for 5 minutes. Cells were then resuspended in pre-warmed expansion medium. Viability and yield were assessed using AO/PI staining and analysed with cell counter (CountStar, Mira FL-S). Recovered cells were then seeded onto Matrigel, and medium was refreshed after 2 days.

### Tissue Sections

Fresh human pancreas samples were cut into small pieces and fixed with 4% PFA at 4□ or 10% neutral formalin at room temperature for 24 hours. After washing for three times with PBS, samples were overlaid with OCT and stored at -80□ for frozen sectioning. The embedded samples were sectioned using a freezing microtome (Leica, 3050S) and store at -80□ until use.

### Immunofluorescence, whole mount staining and microscopy

For frozen section staining, sections were incubated with primary antibodies at 4 °C overnight . After three washes with PBST (PBS+0.1% Triton-X 100), sections were incubated with secondary antibodies and DAPI at room temperature for 2 hours. Sections were then washed three times with PBST and mounted for confocal imaging. For paraffin section staining, after dewaxing and rehydration, a similar procedure of staining was followed.

To perform whole mount staining, human primary islets or cultured organoids were fixed in 4% PFA at 4°C for 1 hour and washed 3 times with PBS. After blocking with a buffer containing 10% FBS in PBST at room temperature for 1 hour , organoids were incubated with primary antibodies diluted in blocking buffer at 4°C overnight. Following three washes with PBST, organoids were incubated with secondary antibodies and DAPI at 4°C overnight. Then organoids were washed, mounted with mounting medium, and imaged using confocal microscopy.

All confocal images were captured by Leica SP8 confocal detection system fitted on a Leica DMI6000 microscope or Olympus FV4000 confocal detection system fitted on a BX63 microscope.

### H&E staining

Tissue sections were deparaffinized in histoclear and rehydrated through a graded ethanol series (100%, 95%, 70%) to distilled water. Sections were stained with hematoxylin for 5-10 minutes, rinsed in tap water, and blued in ammonia water. After rinsing, sections were counterstained with eosin for 1-2 minutes, followed by dehydration through graded ethanol and clearing in histoclear. Finally, sections were mounted with a coverslip using a synthetic resin and allowed to dry before microscopic examination.

### Flow cytometry

For intracellular protein analysis, PICOs were dissociated into single-cell suspensions by incubating with TrypLE^TM^ Express for 5-8 min in a 37 °C incubator. The reaction was termination with PBS containing 5% FBS, followed by gentle manual pipetting to ensure complete dissociation. Cells were filtered through a 70 μm cell strainer, fixed and permeabilized using Fixation/Permeabilization Solution at 4°C for 60 minutes. Cells were washed twice with wash buffer (1x Perm/Wash buffer containing 5% FBS), then incubated with primary antibodies diluted with wash buffer at 4°C overnight. After two washes with wash buffer, cells were incubated with secondary antibodies at 4 °C overnight. Following two final washes, cells were filtered through a 40 μm cell strainer, analyzed using FACS Fortessa cytometer (BD Biosciences). Data were analyzed with FlowJo v.10.9.

### RT-qPCR

RNA isolation of primary cells or organoids were performed following the manufacturer’s protocol. Samples were lysed in RNAiso plus. Purified RNA was reverse- transcribed into cDNA with the PrimeScript RT Master Mix. qRT–PCR was performed using SYBR Green Master Mix, with fluorescence detection performed on an Applied Biosystems QuantStudio^TM^ 6 Flex machine. Data was analyzed using ΔΔCt method. All the qPCR primer sequences are provided in Table S3.

### Glucose-stimulated C-Peptide secretion

For C-peptide ELISA test, PICOs or human primary islets were first washed with Krebs buffer for 3 times and then pre-incubated in low glucose (2 mM) for 2 hours to remove residual insulin. Following 3 washes with Krebs buffer, the organoids were incubated in low-glucose (2 mM) for 30 minutes and the supernatant was collected. Then the organoids were washed 3 times with Krebs buffer, incubated in high glucose (20 mM) for 30 minutes and the supernatant was collected. Secreted C-peptide in the collected supernatants was quantified using human C-peptide ELISA kit according to manufacturer’s instructions.

### Arginine-stimulated glucagon secretion

For glucagon ELISA test, PICOs or human primary islets were first washed with Krebs buffer for 3 times and then incubated with 10 mM glucose for 30 minutes. The supernatant was collected. The organoids were washed 3 times again in Krebs buffer and incubated in 10 mM glucose supplemented with 15 mM arginine for 10 minutes. The supernatant was collected. Secreted glucagon in the collected supernatants was quantified using human glucagon ELISA kit according to manufacturer’s instructions.

### Transmission electron microscope (TEM)

To prepare samples for TEM, PICOs or human primary islets were fixed in 2.5% glutaraldehyde at 4°C overnight. After three washes with 0.1M phosphate buffer, samples were treated with 1% aqueous osmium tetraoxide at room temperature for 1 hour. Samples were sequentially dehydrated in an ethanol series (30%, 50%, 70%, 90% and 100%) for 10 min per concentration, followed by two 15-minute incubations in acetone. Dehydrated samples were infiltrated with mixture of acetone:epoxy resin embedding medium 1:1 at room temperature for 2 hours , then change to 100% epoxy resin embedding medium and incubate overnight. The next day, transfer the samples to embedding molds, and place in a 60 °C oven for 48 hours to allow polymerization. 70 nm ultrathin sections were collected onto 200-mesh copper grids. Sections were stained with 2% uranyl acetate aqueous solution at room temperature for 10 minutes in dark and then couterstained with lead citrate for 5 minutes.

For immunolabeling, PICOs or human primary islets were fixed in a solution containing 4% paraformaldehyde and 0.2% glutaraldehyde, followed by post-fixation with aqueous osmium tetraoxide. After gradual dehydration, samples were infiltrated with LRWhite resin in graded ethanol:resin mixtures (3:1,1:1,1:3), followed by pure resin incubation overnight at 4°C. Samples were transferred to capsules and polymerized at 55°C for 48 hours. Ultrathin sections were cut and collected. Sections were blocked, then incubated with primary antibody. After washing, sections were incubated with gold-conjugated secondary antibody. The stained sections were then fixed in 1% glutaraldehyde and washed. Finally, sections were contrasted with uranyl acetate and lead citrate before examination.

Images were acquired with FEI Tecnai G2 Spirit Transmission electron microscope.

### Ca^2+^ imaging

PICOs were harvested after dispase dissociation and plated in 35mm glass-bottom dish without Matrigel. After 48hours, the organoids were washed twice with Krebs buffer and incubated with Fluo4-AM for 30 minutes. Following incubation, Fluo4-AM was washed out with Krebs buffer, and the organoids were incubated in 37 °C for an additional 15 minutes. The organoids were then imaged using *Nikon AI* inverted confocal microscope to acquire high-resolution time series images at 10-second intervals. The glucose challenge protocol during imaging consisted of the following sequential steps: 5 minutes in low glucose (2 mM), Krebs wash, 5 minutes in high glucose (20 mM), and another Krebs wash. This sequential treatment was repeated twice, followed by a final 5-minute incubation of 2 mM glucose containing 30 mM KCl. For the measurement of the average fluorescence intensity, images were quantified using Imaris software (v9.5.0).

### Mouse transplantation studies

Diabetes was induced in SCID/Beige recipient mice by a single intraperitoneal injection of streptozotocin (STZ, 160 mg/kg in a citrate buffer, pH 4.5) 7 days prior to engraftment. Mice with nonfasted blood glucose levels exceeding 20 mM, as measured by a OneTouch glucometer using tail vein blood, were selected as diabetic recipients. 800∼1000 PICOs or 300 islets were engrafted to the recipient mice. Following engraftment, nonfasted blood glucose and body weight were routinely monitored after engrafting for 1 week. Glucose tolerance tests (GTTs) were performed following standard protocol. In brief, mice were fasted overnight and administered an intraperitoneal injection of glucose (2g/ kg body weight). Blood glucose levels were measured at 0, 15, 30, 60, 90, and 120 min after glucose challenges. Serum samples were collected before and after glucose injection for C-peptide quantification by ELISA. Recipients were sacrificed, and the regenerated tissues were harvested for histological analysis, including staining for endocrine markers.

### Non-human primate transplantation

Cynomolgus monkey was obtained and house in Wincon Theracells Biotechnologies co., LTD. All animal care and handling procedures were performed in accordance with the IACUC guidelines established by the facility. A male Cynomolgus macaque was used as recipient for human islet organoids transplantation.

#### Diabetes induction

Diabetes was induced according to previously report^86^. In brief, a single dose of STZ (90 mg/kg) was injected intravenously after overnight fasting, followed by administration of normal saline (40–50 ml) for hydration. Blood glucose was monitored every hour over the first 12 h after STZ injection and thereafter three times a day. Exogenous insulin injections commenced 2 d after STZ treatment. The levels of blood glucose, C-peptide and HbA1c were recorded before PICO transplantation. *Immunosuppression strategy.* The immunosuppression regimen was started 7 d before transplantation (day 0). The protocol was designed based on previously published studies^87,121^. In brief, rituximab (Rituxan, Roche), ATG (Genzyme Polyclonals), Basiliximab (Simulect, Novartis) were used for immunosuppression induction. Methylprednisolone (Solu-Medrol, Pfizer) were administered 10 min before rituximab and ATG treatments to reduce allergic reactions. Sirolimus (Rapamune, Pfizer) and tacrolimus (Prograf, Astellas) were administered daily for immunosuppression maintenance. The dosages of sirolimus and tacrolimus were adjusted according to trough blood levels (tacrolimus 4∼10 ng/ml; sirolimus 4∼10 ng/ml). Cobra Venom Factor (Quidel) and Etanercept (Enbrel, Pfizer) were used for alleviation of instant blood-mediated inflammatory reaction.

#### Transplant surgeries

The transplantation procedures were performed as previously reported^87^. After the i.v. administration of propofol (0.5 ml/kg, Petsun Therapeutics), monkey was anaesthetized with inhalable isoflurane. Heart rate, temperature, blood oxygenation and blood pressure were monitored in real time during the surgical procedure. PICO were infused into the portal vein through a jejunal vein after laparotomy. To prevent instant thrombosis, heparin sodium (150 IU/kg, i.h., Fluxum, Alfasigma) was injected subcutaneously at 2 h and 8 h postoperatively. Antibiotic treatment was continued for 7 d. Pain relief medication was administered for 3 d. Glucose level, HbA1c, body weight, complete blood count, serum creatinine and liver function analysis were routinely performed.

### Sample preparation and scRNA-seq with 10X genomics

Two donor pancreas samples were processed for primary human pancreatic scRNA-seq. 13,000 single cells were loaded per sample, resulting in 2,119 or 1,249 recovered cells for each donor, respectively. The 2 datasets were then integrated with previously published human pancreatic scRNA-seq datasets (6 datasets from different studies) for further analysis.

To investigate the α and β cell maturation process in vitro, PICOs were sampled at both expansion and maturation stages. Organoid cells were dissociated into single cells using TrypLE^TM^ Express. Live cells were isolated by FACS. The resulting datasets were integrated with primary human islet cells for comparative analysis.

For all of the scRNA-seq experiments, cell quality was assessed under microscope prior to chip loading, ensuring cell viability exceeding 75% and single-cell purity above 90%. Libraries were prepared using the Chromium Single Cell 3’ Library (v2 or v3) according to manufacturer’s instructions (10X Genomics). Briefly, single cells were partitioned into gel beads emulsion in the GemCode instrument with cell lysis and barcoded reverse transcription of RNA, followed by amplification, shearing, and 5’ adaptor and sample index attachment. Libraries were sequenced on an Illumina Hiseq PE150 or X-ten. Filtering, alignment to the hg38 transcriptome and unique molecular identifier (UMI)-collapsing were performed using the Cell Ranger (v2 or v3) pipeline with default mapping arguments (10X Genomics).

### Analysis of scRNA-seq data

Cell Filtering: We employed the Seurat V4 R package^118^ for a comprehensive set of analyses, encompassing cell filtering, normalization, data integration, dimensionality reduction, clustering, differential expression analysis, and visualization. To construct the Seurat object, we carefully selected genes that were expressed in at least 2 cells, along with cells that exhibited mitochondrial count percentages below 20% and had at least 500 detected genes.

Principal Component Analysis (PCA) Selection: Differentially expressed genes were identified using the “vst” method. Subsequently, the top 3000 differentially expressed genes were chosen for PCA. The selection of principal components (PCs) was guided by an elbow plot, and 20 PCs were ultimately utilized for the analysis.

Dimensional Reduction, Cell Clustering, and Data Display: Dimensional reduction was carried out using the Uniform Manifold Approximation and Projection (UMAP) algorithm. Cell clustering was based on the shared-nearest neighbor (SNN) method. Clusters were then grouped into one cell type according to the similarity of their expression profiles and the expression of marker genes. Dot plots, heat maps, individual UMAP plots, and violin plots for the specified genes were generated using the DoHeatmap and FeaturePlot functions of the Seurat toolkit.

Integration of PICO-Expansion, PICO-Maturation, and Primary Human Pancreatic scRNA-seq Datasets: After individually filtering each dataset, we integrated PICO-exp, PICO-maturation, and primary human islet cell datasets, which included six published datasets and two newly generated datasets. This integration was achieved using the FindIntegrationAnchors and IntegrateData functions in Seurat, followed by CellCycleScoring, PCA analysis, and UMAP analysis.

Integration of PICO-Maturation Cells with Developing Endocrine Cells: Developing endocrine cells including BP, EP, alpha, beta, delta, and epsilon cells were extracted from a publicly dataset (GSA: PRJCA010743) of human pancreatic epithelial cells at post-conception weeks (PCW) 7–11, which was originally generated and reported by Zhuo Ma et al.^82^. Subsequently, we repeated the aforementioned analysis process with modified parameters. Briefly, canonical correlation analysis (CCA) was employed during the anchor - finding step. PCA on these variable genes was performed using *RunPCA* (npcs =50), and neighbors were found using principal components via *FindNeighbors*. This was followed by Leiden clustering (*FindClusters*, resolution 3) and UMAP dimensionality reduction (*RunUMAP*, 15 neighbors).

Comparison of PICO and Ductal Organoids: Fresh ductal organoid (cultured for 7 days) datasets published by Heather N Zook et al.^84^ were integrated with our PICO datasets and primary human islet cell datasets. Then, we repeated the above - described analysis process with modified parameters. In brief, during the anchor identification step, the reciprocal PCA (rPCA) workflow was applied, which offers a faster and more conservative integration (dims =1:50, k.anchor =7). PCA on these variable genes was performed using *RunPCA* (npcs =50), followed by cell clustering (*FindClusters*, resolution 0.5) and UMAP dimensionality reduction (*RunUMAP*, 20 neighbors).

### Single-cell gene regulatory network analysis by SCENIC

Transcription factor (TF) enrichment and regulon activity were analyzed using the SCENIC R package^119^. The normalized expression matrix generated by Seurat was used as input data. The SCENIC workflow was performed with default parameters. Briefly, coexpression modules between transcription factors and candidate target genes were inferred using the GENIE3 algorithm. Subsequently, RcisTarget was employed to identify modules where the regulator’s binding motif was significantly enriched among target genes, thereby constructing regulons composed exclusively of direct targets. AUCell was then used to score the activity of each regulon in individual cells, generating a binarized activity matrix. Cell state prediction was performed based on the shared activity of regulatory subnetworks. Finally, differentially activated regulons in each cluster were identified using the Wilcoxon rank-sum test, and only regulons with adjusted p-values<0.05 were retained for further analysis.

### Differential expression and annotation

Differential Expression Analysis: Differential expression analysis was conducted within the Seurat framework using the *FindAllMarkers* function with default parameters. For each cluster, the top 20 significantly differentially expressed genes were scrutinized. These genes were then cross-referenced with marker genes identified from a comprehensive search of the published literature to assign annotations to each cluster. Gene Ontology (GO) Analysis: The list of differential expression genes was utilized for Gene Ontology (GO) analysis via the R package clusterProfiler. Only GO terms with a *p*-value less than 0.05 were retained. Subsequently, the results of the GO analysis were visualized using the ggplot2 package, providing a clear and interpretable representation of the data.

### Cell cycle scoring and pseudo-temporal analysis

Single cells were scored for genes associated with the S phase and G2/M phase. Subsequently, each cell was assigned to one of the cell-cycle phases (S, G2/M, or G1) using the *CellCycleScoring* function in the Seurat package. This process allowed for a detailed assessment of the cell - cycle status of individual cells within the dataset. Pseudo-time and Trajectory Analysis: Pseudo-time and trajectory analysis were carried out on the integrated human pancreatic islet cell organoids (PICOs) and primary human pancreatic datasets using the Monocle 3 R package (version 1.3.5). In the Monocle 3 analysis, the gene expression matrix and the Uniform Manifold Approximation and Projection (UMAP) layout were imported into the CellDataSet object. Since the cell-type information had already been transferred from Seurat, there was no need to repeat the entire Monocle 3 workflow for re-identifying differentially expressed genes for clustering, cell-type identification, and trajectory construction.

#### Trajectory Building

The *learn_graph* function in Monocle 3 was applied to construct trajectories with specific parameters (minimal_branch_len = 4, ncenter = 120). This function facilitated the establishment of meaningful cell-state transitions and developmental trajectories within the dataset.

#### Pseudo-time Measurement

The *order_cells* function was utilized to quantify the pseudo - time of each cell. Cells were manually selected as the root nodes of the trajectory graph, which provided a starting point for the analysis of cell-state transitions and the inference of developmental trajectories over pseudo-time.

### Quantification and statistical analysis

Paired or unpaired two-tailed Student’s t tests were performed when two groups of samples were compared. One-way or two-way ANOVA with Tukey’s or Dunnett’s tests were performed when multiple groups were compared. All the p values were calculated using GraphPad PRISM 10 with the following significance: NS p>0.05; * p<0.05; ** p<0.01; *** p<0.001; **** p<0.0001. Statistical details for each experiment can be found in the figures and the legends.

### Data and code availability

The scRNA-seq datasets generated during this study are available at https://www.biosino.org/node/index, with the accession numbers OEP002072 (primary human tissue) and OEP005302 (PICO).

