## Supplemental Figure 1-9 for "Generation and long-term expansion of human pancreatic islet organoids *in vitro*"

Song et al.

Figure S1. PROCR expression was detected in few acinar cells, which cannont form colony in the culture in current condition, related to Figure 1.

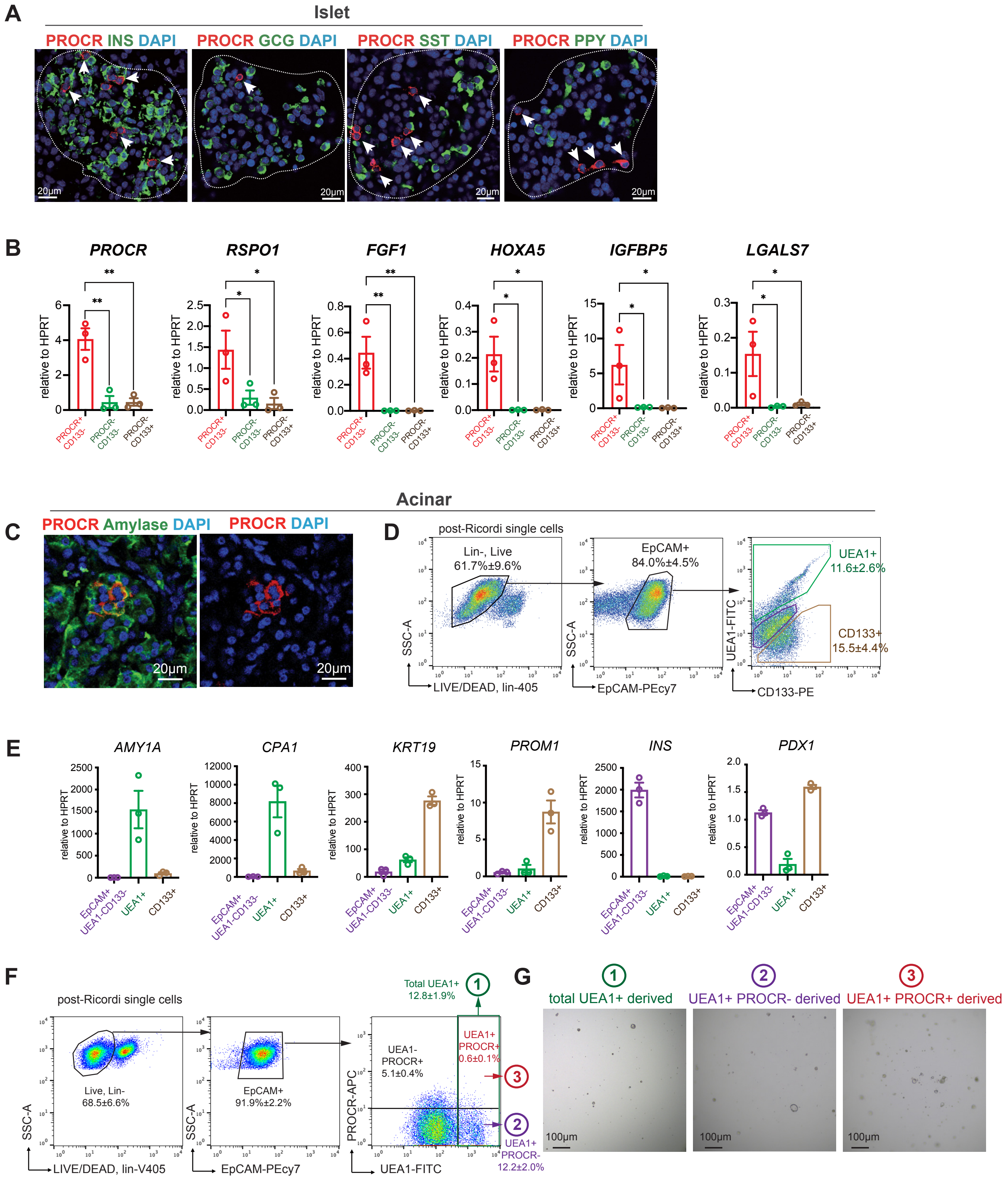

**Figure S1. PROCR expression was detected in few acinar cells, which cannot form colony in the culture in current condition**

(A) Representative confocal images of human pancreatic sections (Donor #2) immunostained for PROCR (red) and endocrine markers (INS, GCG, SST, PPY; green). Dashed lines outline the islet boundary. Arrows denote PROCR+ cells lacking co-localization with β (INS+), α (GCG+), δ (SST+), or PP (PPY+) cells. Scale bar, 20 μm. Experiment repeated with samples from n=3 donors (see Table S1).

(B) RT-qPCR analysis of mouse Procr+ cell signature genes in three FACS-sorted populations (PROCR+, CD133-; PROCR-, CD133-,; PROCR-, CD133+) from human primary islet samples. Data were pooled from n=3 donors (see Table S1), and are represented as mean ± SEM. One-way ANOVA with Tukey's test is used for the comparison of multiple groups. *p<0.05; ** p<0.01. Each symbol represents a donor sample, with data generated from three technical replicates.

(C) Representative confocal images of human pancreatic sections (Donor #2) immunostained for PROCR (red) and acinar marker Amylase (green). Scale bar, 20 μm. Experiment repeated with samples from n=3 donors (see Table S1).

(D) Flow cytometry analysis of cell composition of Ricordi-separated primary human pancreatic cells. The dissociated single cells are stained with Violet Live/Dead dye to exclude dead cells, selected for Lin- (CD45- and CD31-) to exclude blood and endothelial lineage, and selected for EpCAM+ to enrich epithelial cells. UEA1+ (acinar), CD133+ (duct), and UEA1-, CD133- cells were then sorted separately for further analysis. Representative FACS plot is from Donor #12, quantification data were pooled from n=3 donors (see Table S1), and are represented as mean ± SEM.

(E) RT-qPCR validation of the characteristics of three cell populations isolated in (E). Data were pooled from n=3 donors (see Table S1), and are represented as mean ± SEM. Each symbol denotes data from one donor, averaged over three technical replicates.

(F) Flow cytometry analysis of PROCR+ cells and UEA1+ (acinar) cells. Data were pooled from n=3 donors (see Table S1), and are represented as mean ± SEM.

(G) Total UEA1+; UEA1+, PROCR- or UEA1+, PROCR+ cells were FACS sorted and cultured separately. Representative bright-field images (cells derived from Donor #14) at culture day 14 are shown. Scale bar, 100 μm. Experiment also repeated with samples from two other donors (see Table S1).

Song et al.

Figure S2. FACS analysis of human pancreatic PROCR+ islet cells, related to Figure 1.

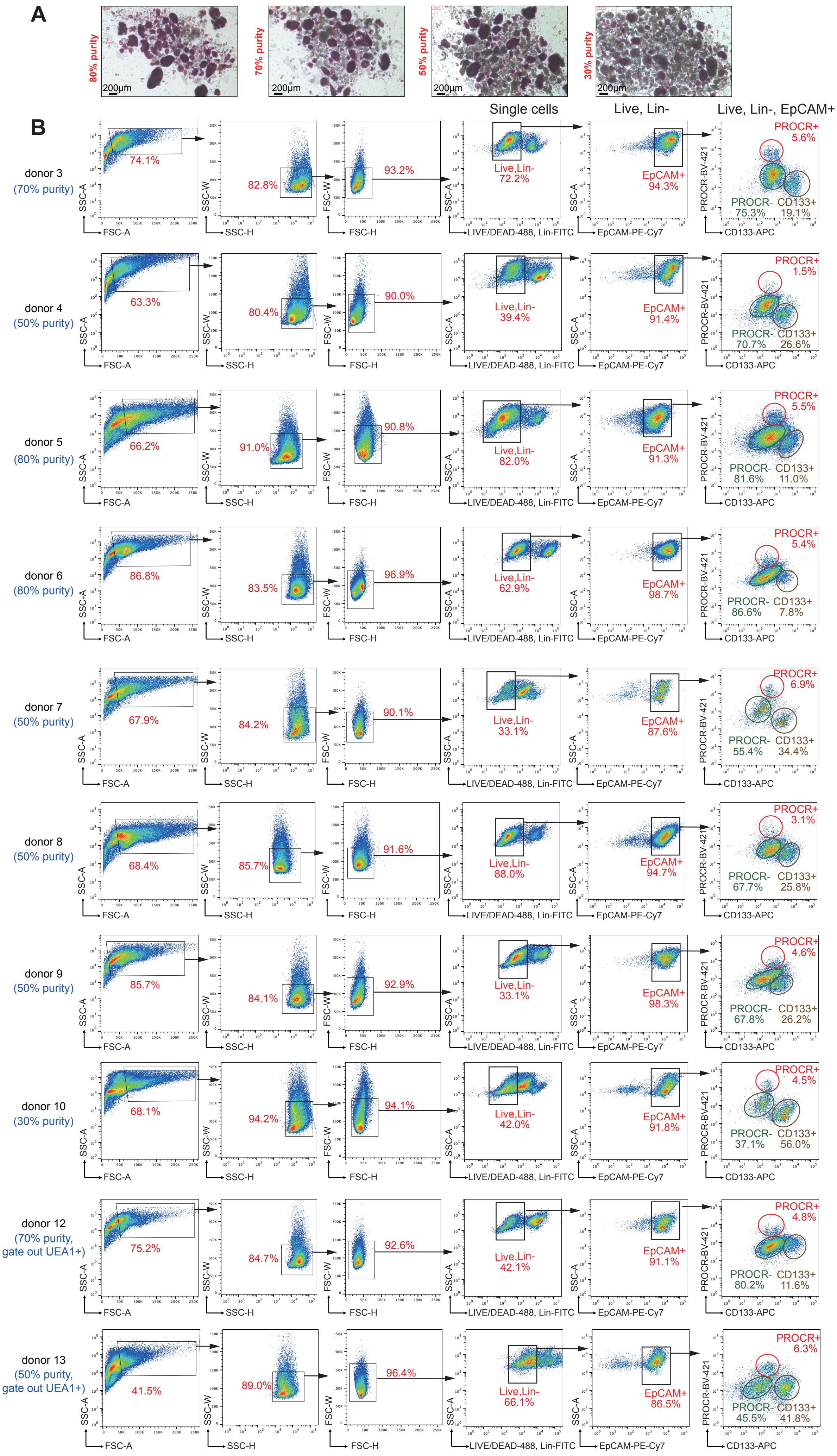

**Figure S2. FACS analysis of human pancreatic PROCR+ islet cells, related to Figure 1.**

(A) Representative bright-field images depict varying purity levels of enriched human islets from Donor #6, as indicated by DTZ staining. Scale bar, 200 μm. The experiment was repeated in all donor samples.
(B) Flow cytometry analysis of PROCR+ cells in primary human pancreatic islet cells (n=10 donors). Dissociated islet single-cell suspensions were stained with Violet Live/Dead dye to exclude dead cells, selected for Lin- (CD45- and CD31-) to exclude blood and endothelial lineage, and enriched for EpCAM+ epithelial cells. CD133 was used to exclude duct cells. For donor #12 and #13, UEA1+ cells are further excluded alongside with CD45+ and CD31+ cells prior to EpCAM+ selection. The percentage of PROCR+ cells in Lin-, EpCAM+ islet cell compartment was analyzed.

Song et al.

Figure S3. Screen for essential factors for human PICO growth, related to Figure 1.

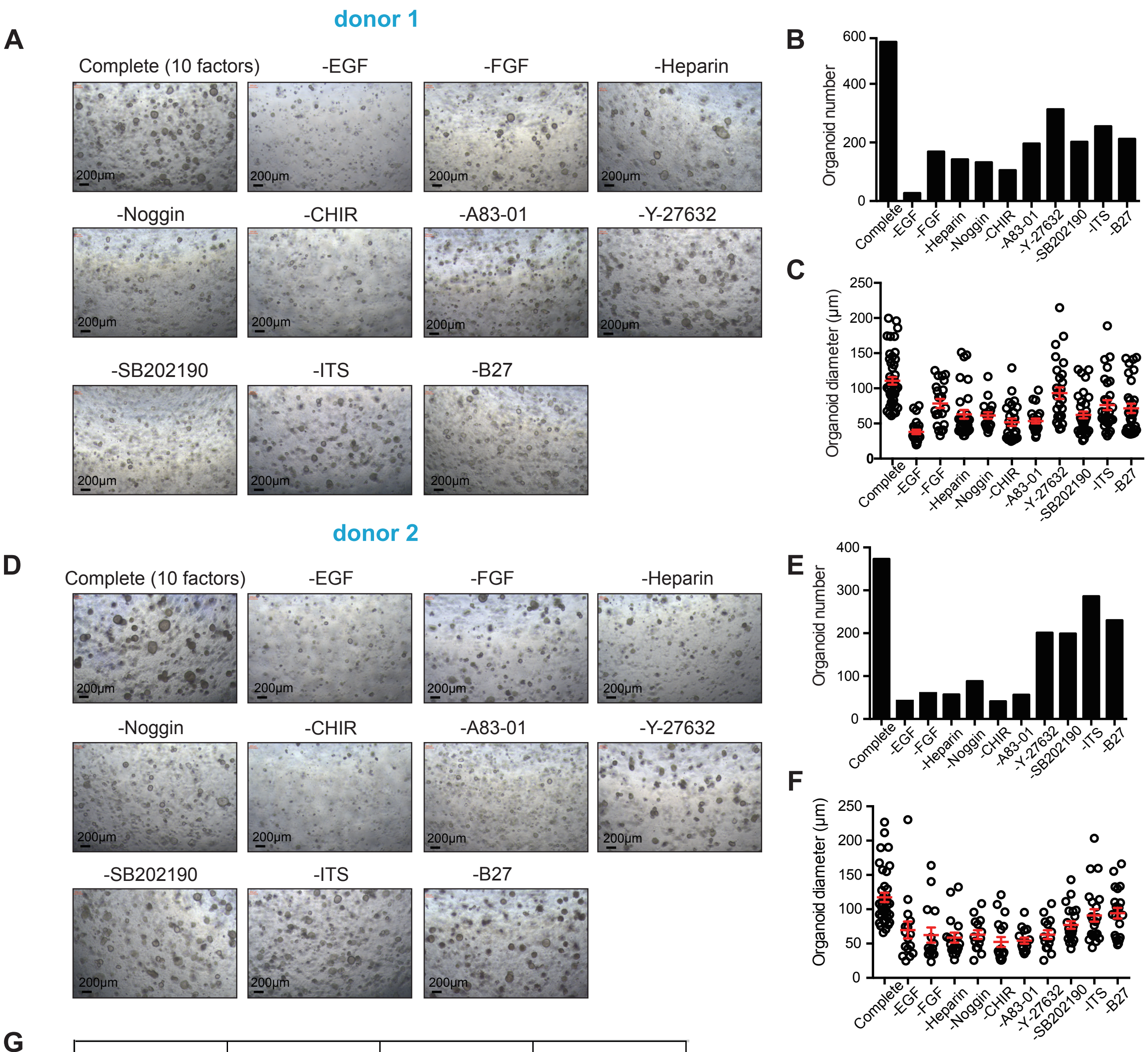


G

| Additives | Concentration | Additives | Concentration |
|---|---|---|---|
| EGF | 50 ng/ml | A83-01 | 0.5 uM |
| FGF | 10 ng/ml | Y27632 | 2.5 uM |
| Heparin | 2 ug/ml | SB202190 | 0.5 uM |
| Noggin | 100 ng/ml | ITS | 1% |
| CHIR | 2.5 uM | B27 | 2% |

**Figure S3. Screen for essential factors for human PICO growth, related to Figure 1.**

(A, D) Representative bright-field images of organoids cultured in different growth factor compositions. Scale bar, 200 μm. The data presented are from 2 independent donors: Donor #1 (A), and Donor #2 (D) (see also Table S1).

(B-C,E-F) Quantification of organoid sizes and numbers under varying growth factor compositions (organoids were initiated from 300,000 unsorted enriched islet cells). Organoid diameters are shown as mean ± SEM (three technical replicates). Data are from 2 independent donors: Donor #1 (B-C), and Donor #2 (E-F) (see also Table S1).

(G) Compositions and concentration of additives in complete expansion medium.

### Figure S4. PICO derived from multiple donors ameliorate diabetes in STZ-induced diabetic mice, related to Figure 2

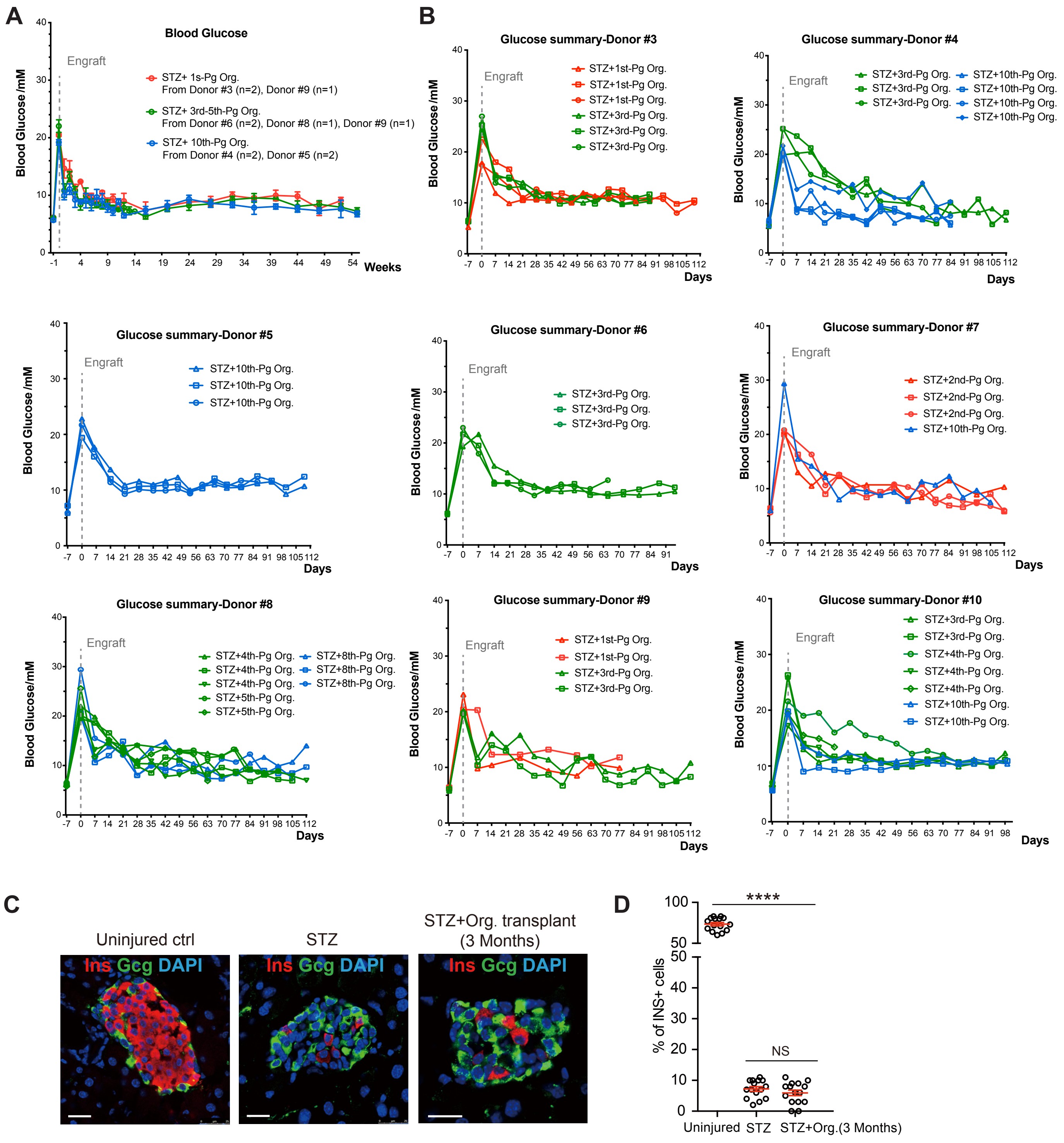

**Figure S4. PICO derived from multiple donors ameliorate diabetes in STZ-induced diabetic mice, related to Figure 2.**

(A-B) SCID-beige mice were induced diabetic with STZ injection 7 days before engrafting. Non-fasting blood glucose levels were monitored. Hyperglycemia was ameliorated in recipients after engrafting PICOs generated from different donors (as indicated), and cultured for various passages (B), for as long as 1 year after transplantation (A). Data from n=8 donors are shown.

(C-D) Representative immunostaining images of INS (β cells) and GCG (α cells) on mice pancreas sections of uninjured ctrl, STZ induced with or without PICOs transplantation. Scale bar, 20 μm.The proportion of INS+ cells from each group is calculated and shown (D). Transplanted organoids were generated from n=3 donors (see Table S1). One-way ANOVA with Tukey's test is used for comparison of multiple groups. NS $p>0.05$; **** $p<0.0001$.

Figure S5. Screen for factors for human PICO maturation, related to Figure 3.

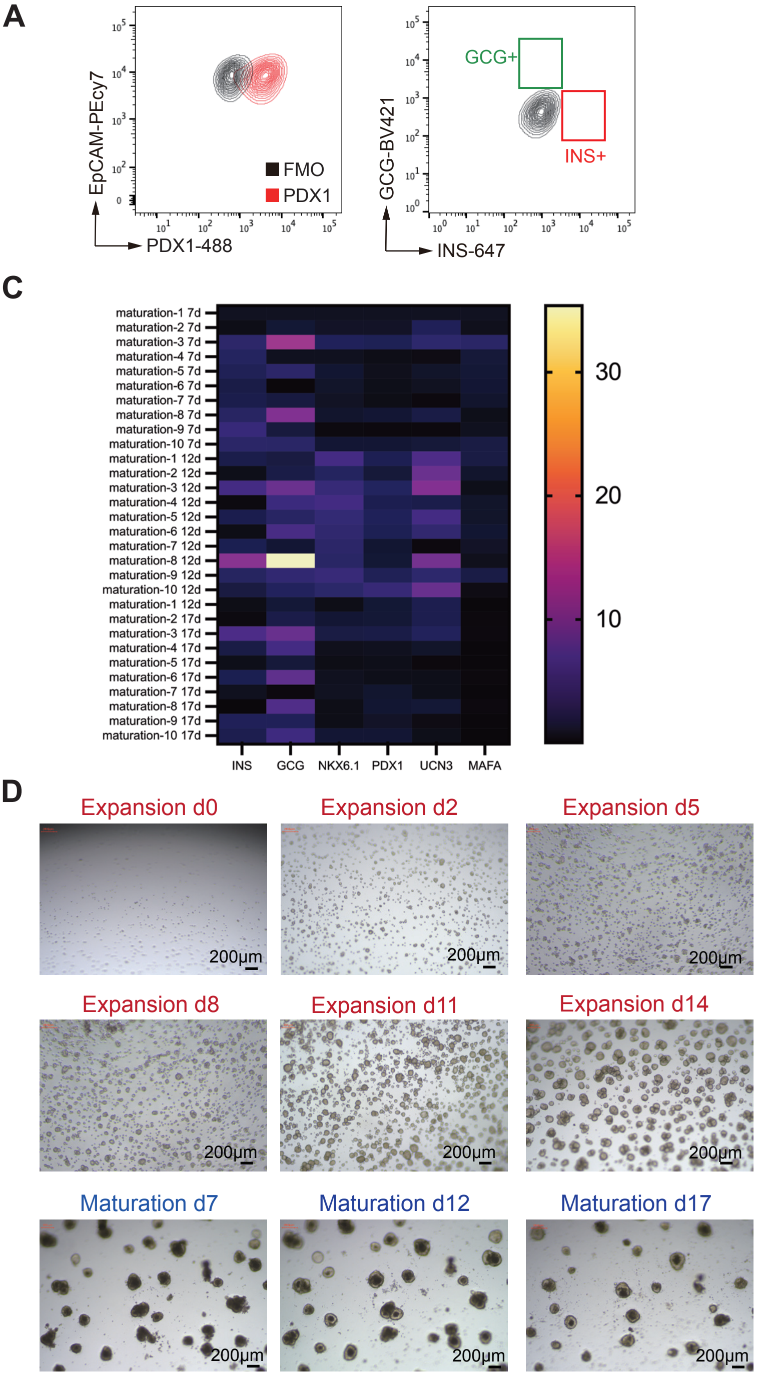


B

| maturation medium | Additives | maturation medium | Additives |
|---|---|---|---|
| 1 | +1% ITS<br>+2% B27<br>+50 ng/ml EGF<br>+0.5 uM A8301<br>+2.5 uM Y27632<br>+100 nM GSIXX | 6 | +1% ITS<br>+2% B27<br>+50 ng/ml EGF<br>+0.5 uM A8301<br>+10 uM Y27632<br>+100 nM GSIXX |
| 2 | +1% ITS<br>+2% B27<br>+50 ng/ml EGF<br>+0.5 uM A8301<br>+2.5 uM Y27632<br>+100 nM GSIXX<br>+1% Glutamax | 7 | +1% ITS<br>+2% B27<br>+0.5 uM A8301<br>+2.5 uM Y27632<br>+100 nM GSIXX |
| 3 | +1% ITS<br>+2% B27<br>+50 ng/ml EGF<br>+0.5 uM A8301<br>+2.5 uM Y27632<br>+100 nM GSIXX<br>+0.25 uM LDN | 8 | +1% ITS<br>+2% B27<br>+0.5 uM A8301<br>+2.5 uM Y27632<br>+100 nM GSIXX<br>+0.25 uM LDN |
| 4 | +1% ITS<br>+2% B27<br>+50 ng/ml EGF<br>+0.5 uM A8301<br>+10 uM Y27632<br>+100 nM GSIXX<br>+0.25 uM LDN | 9 | +1% ITS<br>+2% B27<br>+50 ng/ml EGF<br>+0.5 uM SB431542<br>+2.5 uM Y27632<br>+100 nM GSIXX |
| 5 | +2% B27<br>+50 ng/ml EGF<br>+0.5 uM A8301<br>+2.5 uM Y27632<br>+100 nM GSIXX | 10 | +1% ITS<br>+2% B27<br>+50 ng/ml EGF<br>+0.5 uM A8301<br>+2.5 uM Y27632<br>+100 nM GSIXX<br>+0.25 mM VC |

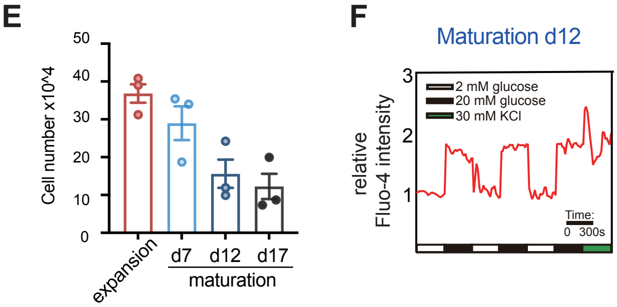


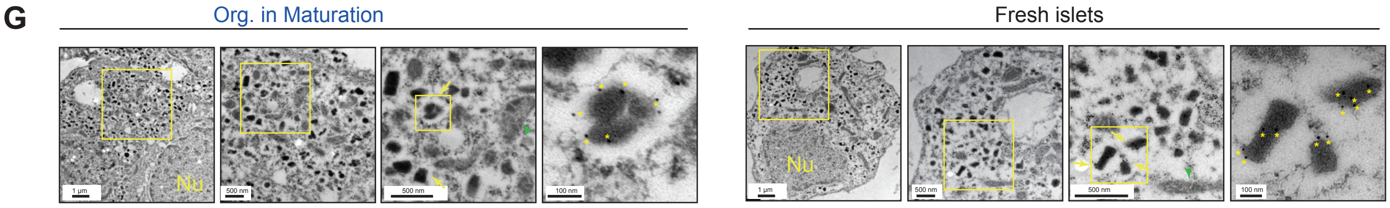

**Figure S5. Screen for factors for human PICO maturation, related to Figure 3.**

(A) Flow cytometry analysis of PDX1 expression (left), INS and GCG expression (right) in PICO under expansion culture conditions. Experiment repeated in n=3 donors (see Table S1).

(B) Compositions and concentrations of different maturation media formulations.

(C) RT-qPCR analysis of endocrine lineage markers (*INS*, *GCG*, *NKX6.1*, *PDX1*, *UCN3* and *MAFA*) expression in PICO cultured in different maturation media and harvested at specified time points. Data are average of three technical replicates. The colors ranging from purple to yellow indicate low to high relative gene expression levels. Experiment repeated in n=3 donors (see Table S1).

(D) Representative bright-field images of PICO formation and maturation (Donor #6). Scale bar, 200 μm.

(E) Quantification of flow cytometry results showing live cell percentage of PICO in expansion medium or after changing to maturation medium over various time periods. Data were pooled from n=3 donors (see Table S1) and are shown as mean ± SEM.

(F) Calcium signaling dynamics in matured organoids subjected to sequential glucose challenges (2, 20, 2, 20, 2, and 20 mM), followed by 30 mM KCl stimulus. The x-axis represents time. Representative data derived from Donor #3. Experiment also repeated with samples from two other donor (see Table S1).

(G) Transmission electron microscopy (TEM) images of granules and immunogold-labeled insulin particles in matured organoid cells (left) and primary islet cells (right). The boxed area is magnified on the right. Yellow arrows indicate INS granules, green arrowsheads indicate mitochondria. Yellow asterisks indicate INS-gold particles. Scale bars, 1 μm; 500 nm; 100 nm. Nu., Nucleus. Experiment repeated in n=3 donors (see Table S1).

Song et al.
Figure S6. Single-cell RNA-seq analysis of PICO and primary islets, related to Figure 4.

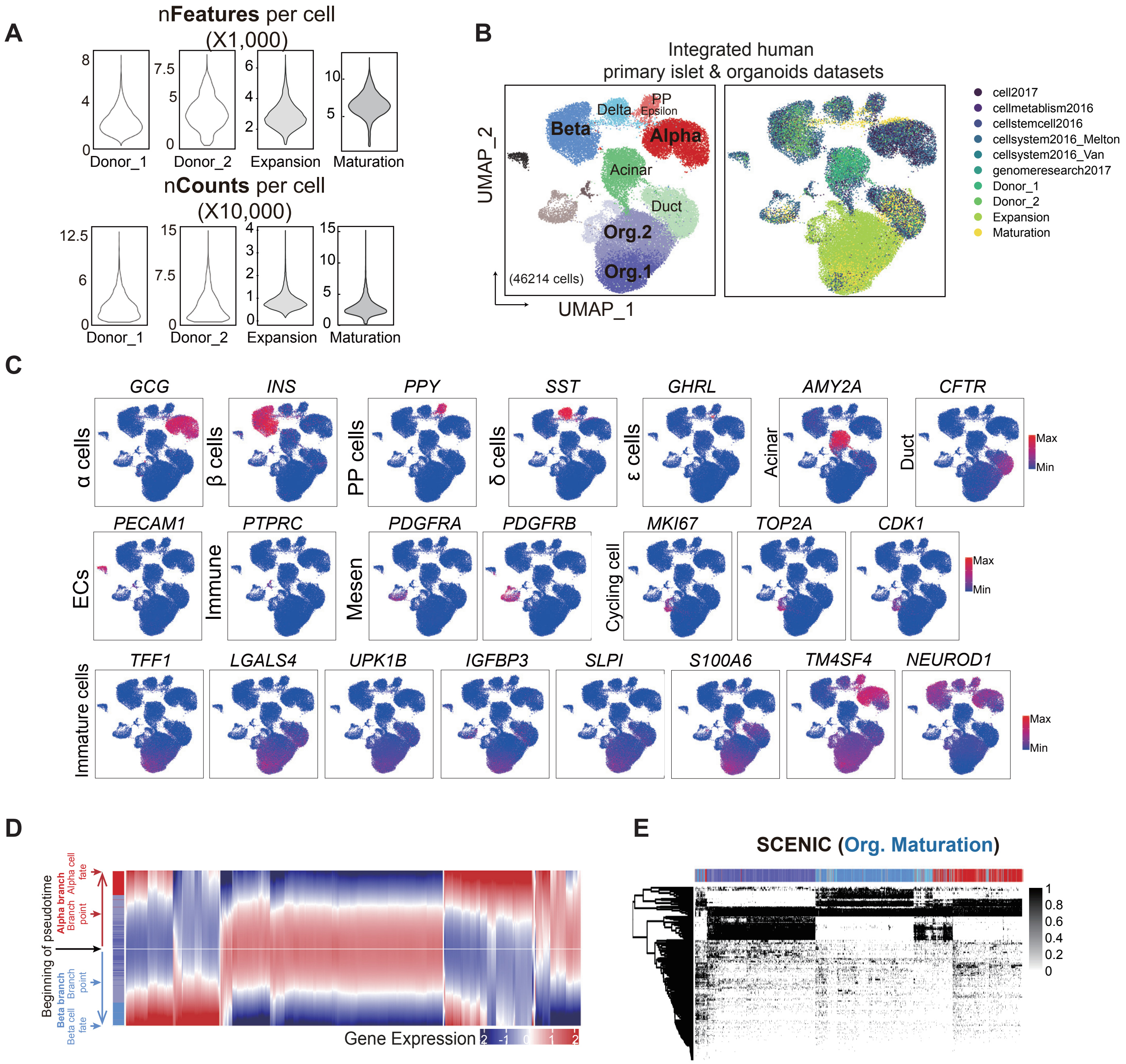

**Figure S6. Single-cell RNA-seq analysis of PICO and primary islets, related to Figure 4.**

(A) Data quality of the hPICO scRNA-seq datasets in expansion (donor #3) and maturation (donor #13) and two primary human pancreatic islet cells (donor #1 and #2) deposited in this study. For expansion-stage dataset, about 15000 viable cells were loaded for sequencing and for maturation-stage dataset, around 10000 viable cells were loaded. Both the number of genes detected per cell (top panel) and the number of UMIs per cell (bottom panel) were shown.

(B) UMAP plot of integrated scRNA-seq data showing the alignment of PICO cells with primary pancreatic cells. The left panel illustrates the distribution of different cell types, including Alpha, Beta, Delta, Epsilon, PP, Acinar, Duct, Org.1, Org.2, Cycling cells, Fibroblast, immune cells and endothelial cells, across 46214 cells. The right panel displays the distribution of each dataset, with colors representing the gradient of pseudotime progression.

(C) Individual gene UMAP plots showing the expression levels ($\log_2$(TPM+1)) of signature genes of each cell cluster.

(D) Gene branched heatmap depicting the expression of genes along each branch in pseudotime. An independent expression pattern is calculated across the entire pseudotime trajectory for each branch. Therefore, the portion of the trajectory before the branch point is displayed for each branch separately. Genes are clustered based on expression pattern across pseudotime.

(E) The heatmap showing the binary activity matrix after applying SCENIC, related to Figure 4E-G.

Song et al.
Figure S7. The molecular similarity between Org. 1 and PROCR+ cells in PICO, related to Figure 4.

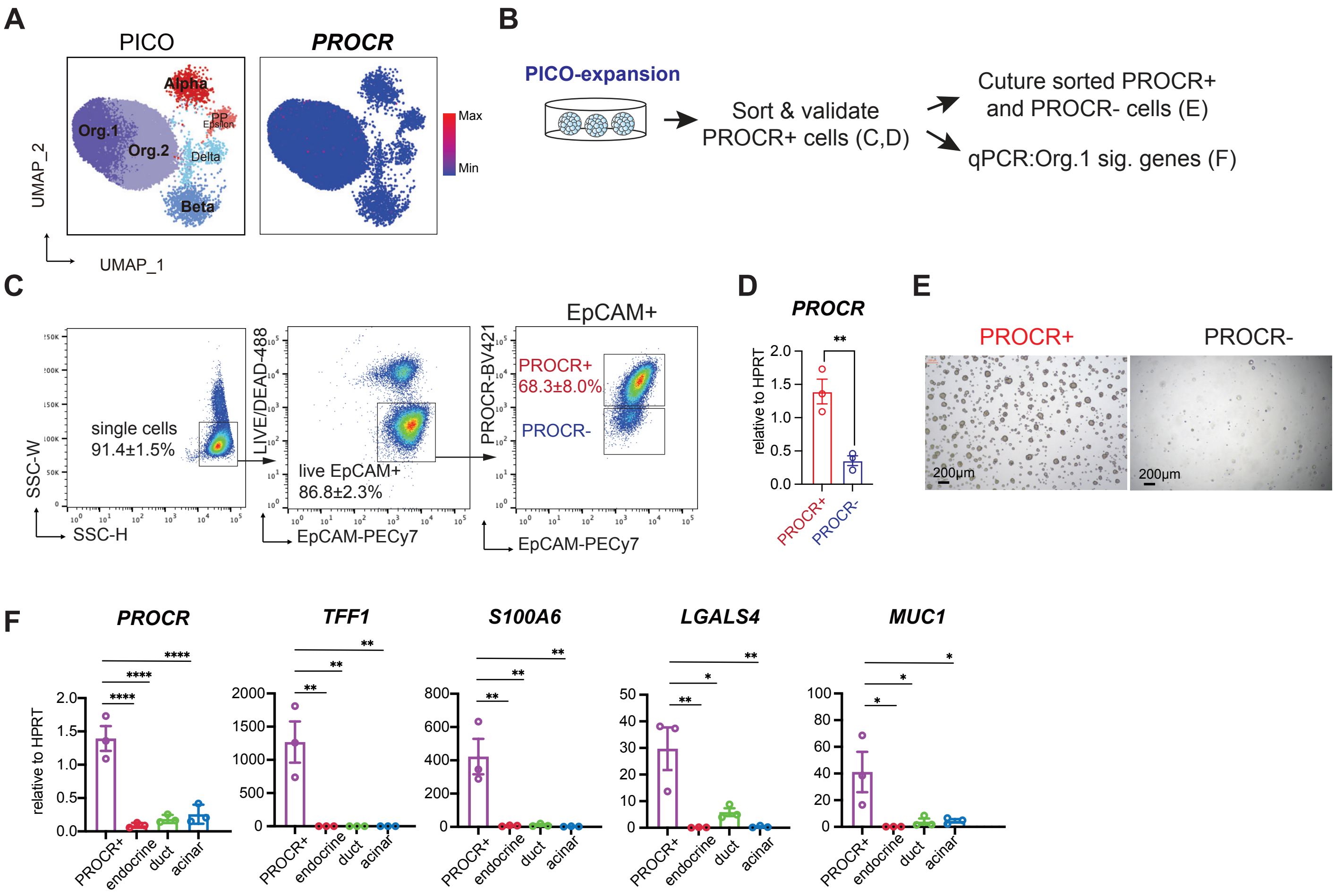

**Figure S7. The molecular similarity between Org. 1 and PROCR+ cells in PICO, related to Figure 4.**

(A) UMAP plots showing the expression levels ($log_2$(TPM+1)) of PROCR in integrated scRNA-seq data.
(B) Illustration of PROCR expression validation in PICO cells during expansion.
(C) Flow cytometry analysis of PROCR expression level in PICO cells. The dissociated single cells were stained with Violet Live/Dead dye to exclude dead cells, EpCAM+, PROCR+ cell population was analyzed. Data were compiled from n=3 donors (see Table S1), and are represented as mean ± SEM.
(D) RT-qPCR validation of *PROCR* expression in FACS sorted PROCR+ and PROCR- cells from cultured PICO under expansion condition. Data were compiled from n=3 donors (see Table S1), and are represented as mean ± SEM. Two-tailed t test is used for comparison. ** p<0.01.
(E) Representative bright-field images of organoids at culture d14 derived from isolated PROCR+ cells and PROCR- cells in PICO (Donor #10). Scale bar, 200 μm. Experiments repeated in n=3 donors (see Table S1).
(F) RT-qPCR analysis of Org.1 signature genes in FACS-isolated PROCR+ cells from PICO, compared with endocrine, duct and acinar cells sorted from fresh pancreatic samples. Data were compiled from n=3 donors (see Table S1), and are represented as mean ± SEM. One-way ANOVA with Tukey's test is used for comparison of multiple groups. * p<0.05; ** p<0.01; **** p<0.0001.

Song et al.

Figure S8. Org.1 exhibited molecular similarity to duct cells, related to Figure 4

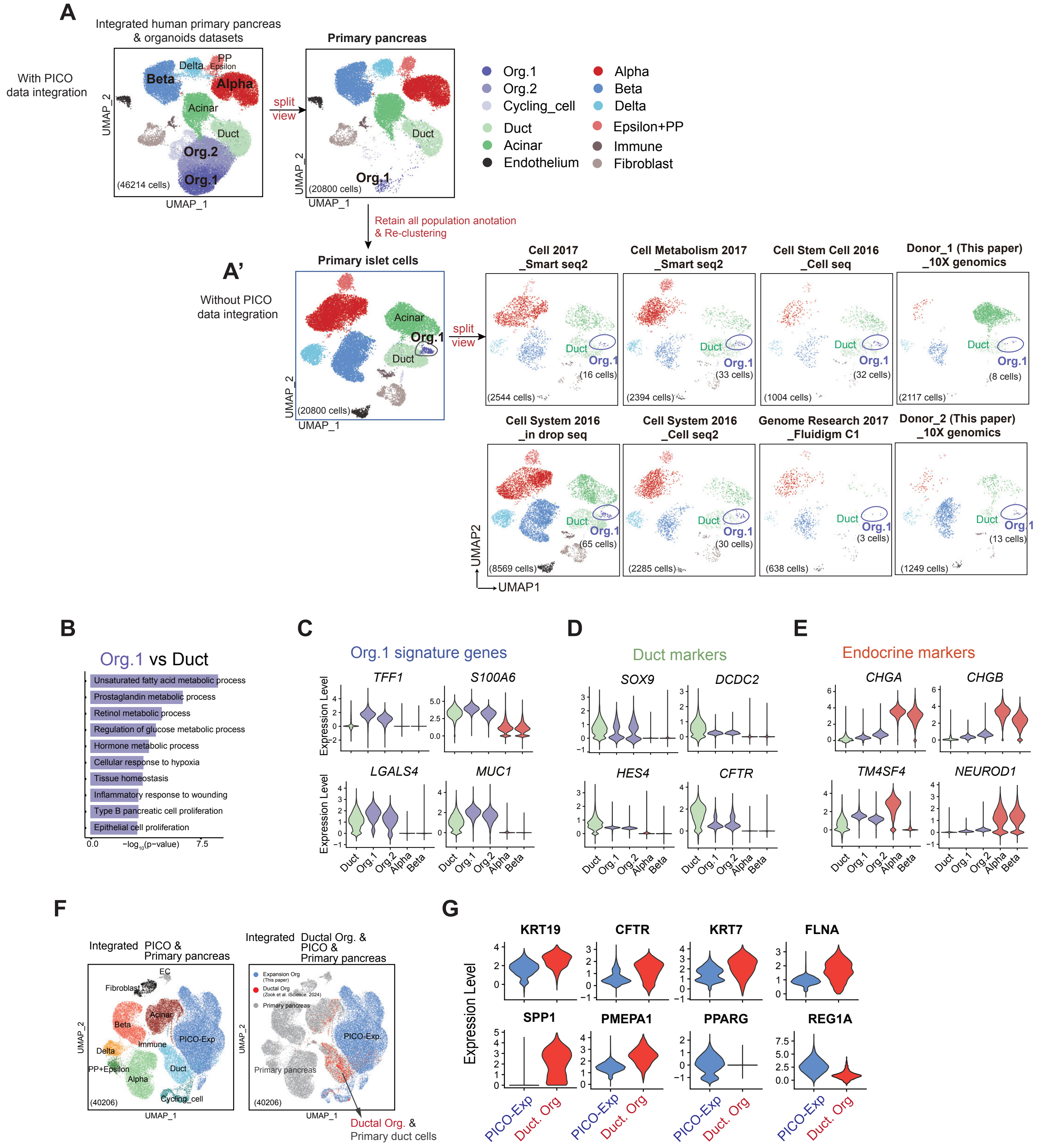

**Figure S8. Org.1 exhibited molecular similarity to duct cells, related to Figure 4.**

(A) Fresh pancreatic data were extracted from integrated datasets, which already incorporated cell-type annotations. UMAP analyses were then performed. After re-clustering, Org.1 cells were found to cluster with ductal cells in the human primary pancreas cell data (A', left panel). split-view UMAP plots of eight human pancreatic scRNA-seq datasets (A', right panel) revealed the presence of rare Org.1 cells across these datasets. Cells were colored according to cluster assignment and annotated post-hoc. Dataset resources and sequencing methods are as indicated. Cell counts of each dataset are labeled in brackets.

(B) Bar plot showing the enriched GO (Gene Ontology) terms of differentially expressed genes in Org.1 cells versus ductal cells. *P*-values were calculated by using *enrichGO* function from R package clusterProfiler with one-sided hypergeometric test.

(C-E) Violin plots showing the expression levels ($\log_2$(TPM+1)) of representative genes of organoid cell (C), ductal (D) and endocrine populations (E).

(F) Human expansion stage PICO single-cell (sc) RNA-seq dataset are integrated with human ductal organoid dataset and primary human pancreatic islet datasets and projected onto UMAP plots, colored by cluster assignment, and annotated post hoc. Both the aligned (left) and split (right) views are shown. Cell counts of each dataset are labeled in brackets.

(G) Violin plots displaying the log2(TPM+1) expression levels of representative genes in ductal organoid cells and expansion stage PICO cells, highlighting the differences between these two cell populations.

### Figure S9. Phenotypic analysis of cynomolgus macaques post transplantation of PICO, related to Figure 5.

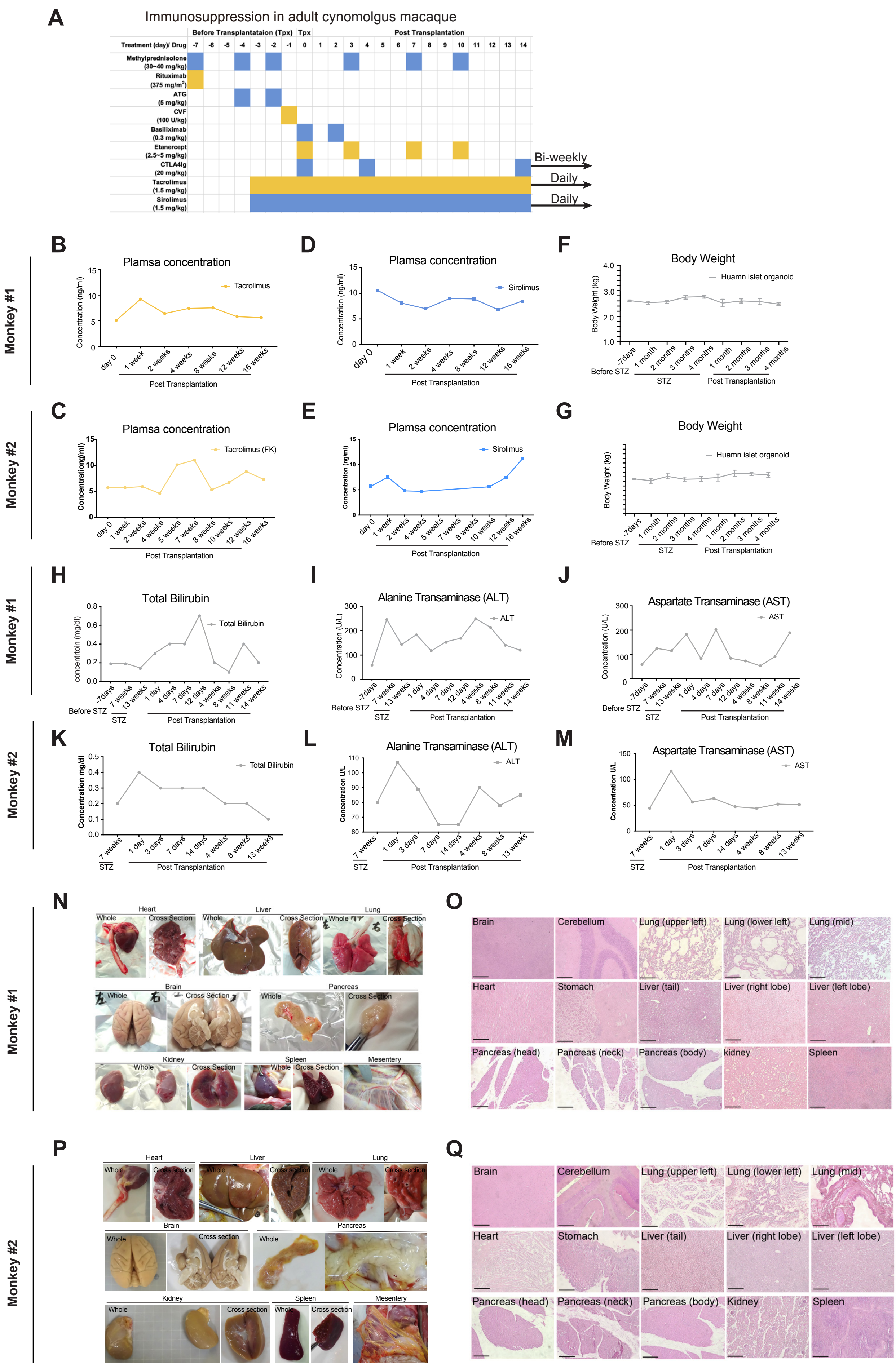

**Figure S9. Phenotypic analysis of *cynomolgus* macaques post transplantation of PICOs, related to Figure 5.**

(A) Schematic illustration describing the overall immunosuppressive regime used.

(B-E) Tacrolimus and Sirolimus plasma concentration of two macaques at various time points post PICO infusion. Day 0 depicts the day of transplantation surgery.

(F, G) Body weight of two macaques throughout experimental stages.

(H-M) Blood biochemistry test results of two macaques at various experimental time points.

(N-Q) Postmortem examination of major organs in transplanted diabetic macaques. Gross anatomy (N, P) and H&E staining (O, Q) of major macaque organs. Scale bar, 250 $\mu$m.
