## Supplemental Table 1-3 for "Generation and long-term expansion of human pancreatic islet organoids *in vitro*"

### Table S1, Donor demographics and related data

| Donor ID | Sample ID | Age of donor (years) | Gender | BMI | Tissue ischaemic time | Underlying pathology | Used in experiments represented in figures |
|---|---|---|---|---|---|---|---|
| #1 | RJ30720 | 41-50 | male | 25 7 | 11hr | none | Figure 2E-F, 2I, 3D, 4A-D,S1A, S2A, S3A-C, S6A-D, S7A, S8A-G |
| #2 | RJ30926 | 41-50 | male | 26 6 | 14hr | none | Figure 3D, 4A-D, S1A, S1C, S2A, S3D-F, S6A-D, S7A, S8A-G |
| #3 | RJ31217 | 51-60 | female | 23 4 | 9hr | none | Figure 1D-H, 2C, 2E-H, 3E-G,4A-B, 4D, S2A-B, S4A-B, S5A, S5C, S5F, S6A-D, S7A, S8A-G,Table S2 |
| #4 | RJ30905 | 41-50 | male | 20 8 | 15hr | none | Figure1D, 1F-H, 2B-C, 2E-I, 3A-C, 3F-H, S2A-B, S4A-B, S5A, S5C-E, Table S2 |
| #5 | RJ31124 | 51-60 | male | 20 2 | 11hr | none | Figure 1D-H, 1L-M, 2C, 2E-I, 3B-C, 3G, 3H, S2A-B, S4A-B, S5F |
| #6 | RJ31129 | 51-60 | male | 20 3 | 7hr | none | Figure 1D-H, 2B-C, 2E, 2G-I, S2A-B, S4A-B, S5D-E |
| #7 | RJ30729 | 51-60 | female | 27 3 | 5hr | none | Figure 1D, 2B-C, 2E, 2G-H, S1A, S1C, S2A-B, S4B-D, S5A, S5C-E |
| #8 | RJ40111 | 51-60 | male | 22 | 7hr | none | Figure 1D, 2C, 2E-J, 3B-F, 3I, S2A-B, S4A-D, S5F-G, Table S2 |
| #9 | RJ40115 | 51-60 | female | 35 2 | 8hr | none | Figure 1D, 2E-I, S1B, S2A-B, S4A-B, S5G |
| #10 | ZH31013 | 51-60 | male | 20 7 | 14hr | none | Figure 1D, 1I-M, 2E-J, S1B, S2A-B, S4B-D, S7C-F |
| #11 | TJ10714 | 31-40 | male | 21 | 8hr | none | Figure 2G-H, 3H, 5B, 5D, 5F, 5H, 5J, S2A, S9B, S9D, S9F, S9H-J, S9N-O |
| #12 | RJ41028 | 61-70 | male | 18 2 | 6.5hr | none | Figure 1B-D, 1F-M, S1B, S1D-G, S2A-B, S7C-F,Table S2 |
| #13 | RJ41122 | 31-40 | male | 24 8 | 6.5hr | none | Figure 1B, 1D, 1F-K, 2C, 4A-I, S1F-G, S2A-B, S5G, S6A-E, S7A, S8A |
| #14 | RJ41213 | 31-40 | male | 20 3 | 7.5hr | none | Figure 1B-C, S1C-G, S2A, S7C-F |
| #15 | RJ50210 | 41-50 | female | 27 1 | 6.5hr | none | Figure 1C, 5C, 5E, 5G, 5I, 5K, S1D-E, S2A, S9C, S9E, S9G, S9K-M, S9P-Q |

**Table S2. PICO at expansion stage can be cryopreserved and maintained for long term.**

| Donor ID | Cell density before freezing | Cell viability | Restoration rate | Length of $N_2$ storage |
|---|---|---|---|---|
| #8 | 5x10^6 /ml | 93% | 80.90% | 30 days |
| #12 | 4.5x10^6 /ml | 94.08% | 82.1% | 30 days |
| #3 | 7.5x10^6 /ml | 88.65% | 81.5% | 2 months |
| #4 | 4.8x10^6/ml | 86.22% | 80.34% | 3 months |

PICO from various donors were cryopreserved for a range of time length before thawing for validation. Upon thawing, cells were assessed for viability and cell count, which enabled the calculation for restoration rate as shown in the table.

**Table S3. The list of gene-specific primers used in RT-qPCR**

| Gene | Forward primer | Reverse primer |
|---|---|---|
| *HPRT* | GCTATAAATTCTTTGCTGACCTGCTG | AATTACTTTTATGTCCCCTGTTGACTGG |
| *INS* | GCAGCCTTTGTGAACCAACA | GGTGTGTAGAAGAAGCCTCGTT |
| *GCG* | TTCTACAGCACACTACCAGAAGA | CTGGGAAGCTGAGAATGATCTG |
| *PDX1* | CGGAACTTTCTATTTAGGATGTGG | AAGATGTGAAGGTCATACTGGCTC |
| *NKX2.2* | TTCCAGAACCACCGCTACAAG | GGGCGTCACCTCCATACCT |
| *NKX6.1* | GGGCTCGTTTGGCCTATTCGTT | CCACTTGGTCCGGCGGTTCT |
| *UCN3* | GTTGAGGCAGCTGAAGATGG | GGAGGGAAGTCCACTCTCG |
| *MAFA* | GCTCTGGAGTTGGCACTTCT | CTTCAGCAAGGAGGAGGTCA |
| *SOX9* | AGCGAACGCACATCAAGAC | CTGTAGGCGATCTGTTGGGG |
| *KRT19* | AGCTAGAGGTGAAGATCCGCGA | GCAGGACAATCCTGGAGTTCTC |
| *PROM1* | CACTACCAAGGACAAGGCGTTC | CAACGCCTCTTTGGTCTCCTTG |
| *PROCR* | GCTCAATGCCTACAACCGCACT | CGAAGTGTAGGAGCGGCTTGTT |
| *AMY1A* | GATAATGGGAGCAACCAAGTGGC | CAGTATGTGCCAGCAGGAAGAC |
| *KRT7* | TGTGGATGCTGCCTACATGAGC | AGCACCACAGATGTGTCGGAGA |
| *CFTR* | GGAGAGCATACCAGCAGTGACT | TTCCAAGGAGCCACAGCACAAC |
| *CD44* | CCAGAAGGAACAGTGGTTTGGC | ACTGTCCTCTGGGCTTGGTGTT |
| *HNF1B* | CCCAGCAAATCTTGTACCAGGC | ACCTCAGTGACCAAGTTGGAGC |
| *CHGA* | GGTTCTTGAGAACCAGAGCAGC | GCTTCACCACTTTTCTCTGCCTC |
| *CHGB* | ACCAGACAGTCCTGACAGAGGA | TAACAGTGCCCACCGCTCCAAT |
| *NEUROD1* | GGTTATGAGACTATCACTGCTCAG | AGAACTGAGACACTCGTCTGTC |
| *RSPO1* | TGCCTGCTCTGACACCAAGGAG | CAGGTTCCTGTTGGCATTCTCC |
| *FGF1* | ATGGCACAGTGGATGGGACAAG | TAAAAGCCCGTCGGTGTCCATG |
| *HOXA5* | AACCCCAGATCTACCCCTGGAT | CAGGGTCTGGTAGCGCGTGTA |
| *IGFBP5* | CGTGCTGTGTACCTGCCCAATT | ACTTGTCCACGCACCAGCAGAT |
| *LGALS7* | TTGGTTCCTCCCAATGCCAGCA | CTCCTTGCTGTTGAAGACCACC |
| *CPA1* | AGTAAGCGTCCAGCCATCTGGA | GTCGAGAATGGCGGTGAAAGCT |
| *PDX1* | CGGAACTTTCTATTTAGGATGTGG | AAGATGTGAAGGTCATACTGGCTC |
| *TFF1* | CCAGTGTGCAAATAAGGGCTGC | AGGCAGATCCCTGCAGAAGTGT |
| *S1006A* | GGGAGGGTGACAAGCACAC | AGCTTCGAGCCAATGGTGAG |
| *LGALS4* | GGAACAGCCTTCTGAATGGCTC | CCATTGGCGTAAACCTTGAAGCG |
| *MUC1* | CCTACCATCCTATGAGCGAGTAC | GCTGGGTTTGTGTAAGAGAGGC |
| *PAX6* | CTGAGGAATCAGAGAAGACAGGC | ATGGAGCCAGATGTGAAGGAGG |
